# Time-Consistent Reconciliation Maps and Forbidden Time Travel

**DOI:** 10.1101/201053

**Authors:** Nikolai Nøjgaard, Manuela Geiß, Peter F. Stadler, Daniel Merkle, Nicolas Wieseke, Marc Hellmuth

**Affiliations:** Dpt. of Mathematics and Computer Science, University of Greifswald, Walther-Rathenau-Strasse 47, D-17487 Greifswald, Germany; Saarland University, Center for Bioinformatics, Building E 2.1, P.O. Box 151150, D-66041 Saarbrücken, Germany; Bioinformatics Group, Department of Computer Science; and Interdisciplinary Center of Bioinformatics, University of Leipzig, Härtelstraße 16-18, D-04107 Leipzig; Department of Mathematics and Computer Science, University of Southern Denmark, Denmark; Max-Planck-Institute for Mathematics in the Sciences,, Inselstraße 22, D-04103 Leipzig; Inst. f. Theoretical Chemistry, University of Vienna, Währingerstraße 17, A-1090 Wien, Austria; Santa Fe Institute, 1399 Hyde Park Rd., Santa Fe, USA; Parallel Computing and Complex Systems Group, Department of Computer Science, Leipzig University, Augustusplatz 10, 04109, Leipzig, Germany

**Keywords:** Tree Reconciliation, Horizontal Gene Transfer, Reconciliation Map, Time-Consistency, History of gene families

## Abstract

**Background:** In the absence of horizontal gene transfer it is possible to reconstruct the history of gene families from empirically determined orthology relations, which are equivalent to *event-labeled* gene trees. Knowledge of the event labels considerably simplifies the problem of reconciling a gene tree *T* with a species trees *S*, relative to the reconciliation problem without prior knowledge of the event types. It is well-known that optimal reconciliations in the unlabeled case may violate time-consistency and thus are not biologically feasible. Here we investigate the mathematical structure of the event labeled reconciliation problem with horizontal transfer.

**Results:** We investigate the issue of time-consistency for the event-labeled version of the reconciliation problem, provide a convenient axiomatic framework, and derive a complete characterization of time-consistent reconciliations. This characterization depends on certain weak conditions on the event-labeled gene trees that reflect conditions under which evolutionary events are observable at least in principle. We give an 𝒪(|*V*(*T*)|log(|*V*(*S*)|))-time algorithm to decide whether a time-consistent reconciliation map exists. It does not require the construction of explicit timing maps, but relies entirely on the comparably easy task of checking whether a small auxiliary graph is acyclic. The algorithms are implemented in C++ using the boost graph library and are freely available at https://github.com/Nojgaard/tc-recon.

**Significance:** The combinatorial characterization of time consistency and thus biologically feasible reconciliation is an important step towards the inference of gene family histories with horizontal transfer from orthology data, i.e., without presupposed gene and species trees. The fast algorithm to decide time consistency is useful in a broader context because it constitutes an attractive component for all tools that address tree reconciliation problems.

## 1 Background

Modern molecular biology describes the evolution of species in terms of the evolution of the genes that collectively form an organism’s genome. In this picture, genes are viewed as atomic units whose evolutionary history *by definition* forms a tree. The phylogeny of species also forms a tree. This species tree is either interpreted as a consensus of the gene trees or it is inferred from other data. An interesting formal manner to define a species tree independent of genes and genetic data is discussed e.g. in [15].

In this contribution, we assume that gene and species trees are given independently of each other. The relationship between gene and species evolution is therefore given by a reconciliation map that describes how the gene tree is embedded in the species tree: after all, genes reside in organisms, and thus at each point in time can be assigned to a species.

From a formal point of view, a reconciliation map *μ* identifies vertices of a gene tree with vertices and edges in the species tree in such a way that (partial) ancestor relations given by the genes are preserved by *μ*. Vertices in the species tree correspond to speciation events. By definition, in a speciation event all genes are faithfully transmitted from the parent species into both (all) daughter species. Some of the vertices in the gene tree therefore correspond to speciation events. In gene duplications, two copies of a gene are formed from a single ancestral gene and then keep residing in the same species. In horizontal gene transfer (HGT) events, the original remains within the parental species, while the offspring copy “jumps” into a different branch of the species tree. Given a gene tree with event types assigned to its interior vertices, it is customary to define pairwise relations between genes depending on the event type of their last common ancestor [16, 20, 22].

Most of the literature on this topic assumes that both the gene tree and the species tree are known but no information is available of the type of events [17, 35, 44, 42]. The aim is then to find a mapping of the gene tree *T* into the species tree *S* and, at least implicitly, an event-labeling on the vertices of the gene tree *T*. Here we take a different point of view and assume that *T* and the types of evolutionary events on *T* are known. This setting has ample practical relevance because event-labeled gene trees can be derived from the pairwise orthology relation [24, 22]. These relations in turn can be estimated directly from sequence data using a variety of algorithmic approaches that are based on the pairwise best match criterion and hence do not require any *a priori* knowledge of the topology of either the gene tree or the species tree, see e.g. [38,3,32, 2].

Genes that share a common origin (homologs) can be classified into orthologs, paralogs, and xenologs depending whether they originated by a speciation, duplication or horizontal gene transfer (HGT) event [16, 22]. Recent advances in mathematical phylogenetics [19, 24] have shown that the knowledge of these event-relations (orthologs, paralogs and xenologs) suffices to construct event-labeled gene trees and, in some case, also a species tree [18, 25, 20].

Conceptually, both the gene tree and species tree are associated with a timing of each event. Reconciliation maps must preserve this timing information because there are *biologically infeasible* event labeled gene trees that cannot be reconciled with any species tree. In the absence of HGT, biologically feasibility can be characterized in terms of certain triples (rooted binary trees on three leaves) that are displayed by the gene trees [25]. In the presence of HGT such triples give at least necessary conditions for a gene tree being biologically feasible [18]. In particular, the timing information must be taken into account explicitly in the presence of HGT. That is, gene trees with HGT that must be mapped to species trees only in such a way that some genes do not travel back in time.

There have been several attempts in the literature to handle this issue, see e.g. [14] for a review. In [33, 7] a *single* HGT adds timing constraints to a time map for a reconciliation to be found. Time-consistency is then defined as the existence of a topological order of the digraph reflecting all the time constraints. In [40] NP-hardness was shown for finding a parsimonious time-consistent reconciliation based on a definition for time-consistency that in essence considers *pairs* of HGTs. However, the latter definitions are explicitly designed for *binary* gene trees and do not apply to non-binary gene trees, which are used here to model incomplete knowledge of the exact gene phylogenies. Different algorithmic approaches for tackling time-consistency exist [14] such as the inclusion of “time-zones” known for specific evolutionary events. It is worth noting that *a posteriori* modifications of time-inconsistent solutions will in general violate parsimony [33]. So far, no results have become available to determine the *existence* of time-consistent reconciliation maps given the (undated) species tree and the event-labeled gene tree.

Here, we introduce an axiomatic framework for time-consistent reconciliation maps and characterize for given event-labeled gene trees *T* and species trees *S* whether there exists a time-consistent reconciliation map. We provide an 𝒪(|*V*(*T*)|log(|*V*(*S*)|))-time algorithm that constructs a time-consistent reconciliation map if one exists.

## 2 Notation and Preliminaries

We consider *rooted trees T* = (*V*, *E*) (on *L*_*T*_) with root ***ρ***_*T*_ ∈ *V* and leaf set *L*_*T*_ ⊆ *V*. A vertex *v* ∈ *V* is called a *descendant* of *u* ∈ *V*, *v* ⪯_*T*_ *u*, and *u* is an *ancestor* of *v*, *u* ⪰_*T*_ *v*, if *u* lies on the path from ***ρ***_*T*_ to *v*. As usual, we write *v* ≺_*T*_ *u* and *u* ≻_*T*_ *v* to mean *v* ⪯_*T*_ *u* and *u* ≠ *v*. The partial order ⪰_*T*_ is known as the *ancestor order* of *T*; the root is the unique maximal element w.r.t ⪰_*T*_. If *u* ⪯_*T*_ *v* or *v* ⪯_*T*_ *u* then *u* and *v* are *comparable* and otherwise, *incomparable*. We consider edges of rooted trees to be directed away from the root, that is, the notation for edges (*u*, *v*) of a tree is chosen such that *u* ≻_*T*_ *v*. If (*u*, *v*) is an edge in *T*, then *u* is called *parent* of *v* and *v child* of *u*. It will be convenient for the discussion below to extend the ancestor relation ⪯_*T*_ on *V* to the union of the edge and vertex sets of *T*. More precisely, for the edge *e* = (*u*, *v*) ∈ *E* we put *x* ≺_*T*_ *e* if and only if *x* ⪯_*T*_ *v* and *e* ≺_*T*_ *x* if and only if *u* ⪯_*T*_ *x*. For edges *e* = (*u*, *v*) and *f* = (*a*, *b*) in *T* we put *e* ⪯_*T*_ *f* if and only if *v* ⪯_*T*_ *b*. For *x* ∈ *V*, we write *L*_*T*_ (*x*) := {*y* ∈ *L*_*T*_ | *y* _*T*_ *x*} for the set of leaves in the subtree *T* (*x*) of *T* rooted in *x*.

For a non-empty subset of leaves *A* ⊆ *L*, we define lca_*T*_(*A*), or the *least common ancestor of A*, to be the unique ⪯_*T*_-minimal vertex of *T* that is an ancestor of every vertex in *A*. In case *A* = {*u*, *v*}, we put lca_*T*_(*u*, *v*) := lca_*T*_({*u*, *v*}). We have in particular *u* = lca_*T*_(*L*_*T*_(*u*)) for all *u* ∈ *V*. We will also frequently use that for any two non-empty vertex sets *A*, *B* of a tree, it holds that lca(*A* ⋃ *B*) = lca(lca(*A*), lca(*B*)).

A *phylogenetic tree* is a rooted tree such that no interior vertex in *v* ∈ *V* \ *L*_*T*_ has degree two, except possibly the root. If *L*_*T*_ corresponds to a *set of genes* 𝔾 or *species* 𝕊, we call a phylogenetic tree on *L*_*T*_ *gene tree* or *species tree*, respectively. In this contribution we will **not** restrict the gene or species trees to be binary, although this assumption is made implicitly or explicitly in much of the literature on the topic. The more general setting allows us to model incomplete knowledge of the exact gene or species phylogenies. Of course, all mathematical results proved here also hold for the special case of binary phylogenetic trees.

In our setting a gene tree *T* = (*V*, *E*) on 𝔾 is equipped with an *event-labeling* map *t*: *V* ⋃ *E* ⟶ *I*⋃{0,1} with *I* = {•, ◻, △, ⨀} that assigns to each interior vertex *v* of *T* a value *t* (*v*) ∈ *I* indicating whether *v* is a speciation event (•), duplication event (◻) or HGT event (△). It is convenient to use the special label ⨀ for the leaves *x* of *T*. Moreover, to each edge *e* a value *t*(*e*) ∈ {0,1} is added that indicates whether *e* is a *transfer edge* (1) or not (0). Note, only edges (*x*,*y*) for which *t* (*x*) = △ might be labeled as transfer edge. We write ε = {*e* ∈ *E* | *t*(*e*) = 1} for the set of transfer edges in *T*. We assume here that all edges labeled “0” transmit the genetic material vertically, that is, from an ancestral species to its descendants.

We remark that the restriction *t*_|*V*_ of *t* to the vertex set *V* coincides with the “symbolic dating maps” introduced in [6]; these have a close relationship with cographs [19, 21, 23]. Furthermore, there is a map **σ**: 𝔾 → 𝕊 that assigns to each gene the species in which it resides. The set **σ**(*M*), *M* ⊆ 𝔾, is the set of species from which the genes *M* are taken. We write (*T*; *t*, **σ**) for the gene tree *T* = (*V*, *E*) with event-labeling *t* and corresponding map **σ**.

Removal of the transfer edges from (*T*; *t*, **σ**) yields a forest 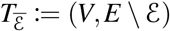 that inherits the ancestor order on its connected components, i.e., 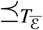 iff *x* ⪯_*T*_ *y* and *x*,*y* are in same subtree of 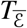 [40]. Clearly 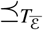 uniquely defines a root for each subtree and the set of descendant leaf nodes 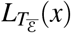.

In order to account for duplication events that occurred before the first speciation event, we need to add an extra vertex and an extra edge “above” the last common ancestor of all species in the species tree *S* = (*V, E*). Hence, we add an additional vertex to *V* (that is now the new root ***ρ**s* of *S*) and the additional edge (***ρ***_*S*_, lca_*S*_(𝕊)) to *E*. Strictly speaking *S* is not a phylogenetic tree in the usual sense, however, it will be convenient to work with these augmented trees. For simplicity, we omit drawing the augmenting edge (***ρ**s*, lca_*S*_(𝕊)) in our examples.

## 3 Observable Scenarios

The true history of a gene family, as it is considered here, is an arbitrary sequence of speciation, duplication, HGT, and gene loss events. The applications we envision for the theory developed, here, however assume that the gene tree and its event labels are inferred from (sequence) data, i. e., (*T*;*t*, **σ**) is restricted to those labeled trees that can be constructed at least in principle from observable data. The issue here are gene losses that may completely eradicate the information on parts of the history. Specifically, we require that (*T*;*t*, **σ**) satisfies the following three conditions:

**(O1)** Every internal vertex *v* has degree at least 3, except possibly the root which has degree at least 2.
**(O2)** Every HGT node has at least one transfer edge, *t* (*e*) = 1, and at least one non-transfer edge, *t* (*e*) = 0;
**(O3)**

**(*a*)** If *x* is a speciation vertex, then there are at least two distinct children *v*, *w* of *x* such that the species *V* and *W* that contain *v* and *w*, resp., are incomparable in *S*.
**(*b*)** If (*v*, *w*) is a transfer edge in *T*, then the species *V* and *W* that contain *v* and *w*, resp., are incomparable in *S*.

Condition **(O1)** ensures that every event leaves a historical trace in the sense that there are at least two children that have survived in at least two of its subtrees. If this were not the case, no evidence would be left for all but one descendant tree, i.e., we would have no evidence that event v ever happened. We note that this condition was used e.g. in [25] for scenarios without HGT. Condition **(O2)** ensures that for an HGT event a historical trace remains of both the transferred and the non-transferred copy. If there is no transfer edge, we have no evidence to classify v as a HGT node. Conversely, if all edges were transfers, no evidence of the lineage of origin would be available and any reasonable inference of the gene tree from data would assume that the gene family was vertically transmitted in at least one of the lineages in which it is observed. In particular, Condition **(O2)** implies that for each internal vertex there is a path consisting entirely of non-transfer edges to some leaf. This excludes in particular scenarios in which a gene is transferred to a different “host” and later reverts back to descendants of the original lineage without any surviving offspring in the intermittent host lineage. Furthermore, a speciation vertex *x* cannot be observed from data if it does not “separate” lineages, that is, there are two leaf descendants of distinct children of *x* that are in distinct species. However, here we only assume to have the weaker Condition (O3.a) which ensures that any “observable” speciation vertex *x* separates at least locally two lineages. In other words, if all children of *x* would be contained in species that are comparable in *S* or, equivalently, in the same lineage of *S*, then there is no clear historical trace that justifies *x* to be a speciation vertex. In particular, most-likely there are two leaf descendants of distinct children of *x* that are in the same species even if only 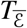 is considered. Hence, *x* would rather be classified as a duplication than as a speciation upon inference of the event labels from actual data. Analogously, if (*v*, *w*) ∈ ε then *v* signifies the transfer event itself but w refers to the next (visible) event in the gene tree *T*. Given that (*v*, *w*) is a HGT-edge in the observable part, in a “true history” *v* is contained in a species *V* that transmits its genetic material (maybe along a path of transfers) to a contemporary species *Z* that is an ancestor of the species *W* containing *w*.

**Figure 1:**
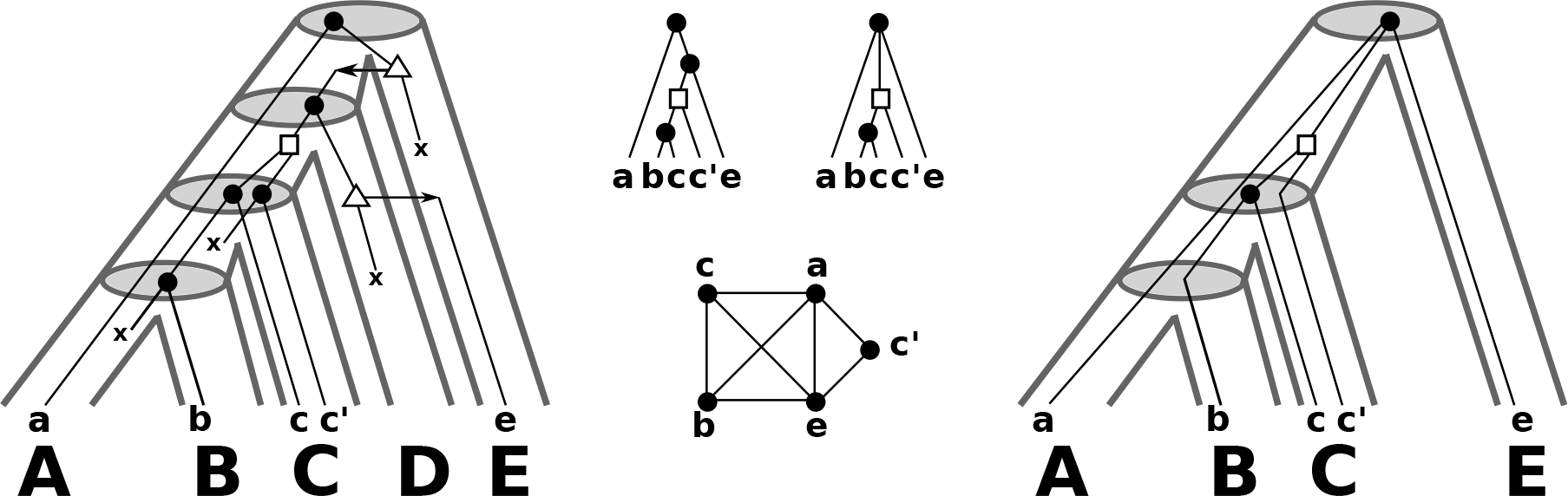
*Left:* A “true” evolutionary scenario for a gene tree with leaf set 𝔾 evolving along the tubelike species trees is shown. The symbol “x” denotes losses. All speciations along the path from the root ***ρ***_*T*_ to the leaf *a* are followed by losses and we omit drawing them. *Middle:* The observable gene tree is shown in the upper-left. The orthology graph *G* = (𝔾, *E*) (edges are placed between genes *x*,*y* for which *t*(lca(*x*,*y*)) = •) is drawn in the lower part. This graph is a cograph and the corresponding *non-binary* gene tree *T* on 𝔾 that can be constructed from such data is given in the upper-right part (cf. [19, 20, 22] for further details). *Right:* Shown is species trees *S* on 𝕊 = **σ**(𝔾) with reconciled gene tree *T*. The reconciliation map *μ*, for *T* and *S* is given implicitly by drawing the gene tree *T* within *S*. Note, this reconciliation is not consistent with DTL-scenarios [40, 5]. A DTL-scenario would require that the duplication vertex and the leaf *a* are incomparable in *S*.

Clearly, the latter allows to have *V* ⪰_*S*_ *W* which happens if the path of transfers points back to the descendant lineage of *V* in *S*. In this case the transfer edge (*v*, *w*) must be placed in the species tree such that *μ*(*v*) and *μ*(*w*) are comparable in *S*. However, then there is no evidence that this transfer ever happened, and thus *v* would be rather classified as speciation or duplication vertex.

Assuming that **(O2)** is satisfied, we obtain the following useful result:

#### Lemma 1.

*Let* 𝒯_1_,…, 𝒯_*k*_ *be the connected components of* 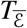 *with roots **ρ***_1_,…, ***ρ***_*k*_, *respectively. If* **(O2)** *holds, then*, 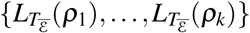 *forms a partition of* 𝔾.

*Proof*. Since 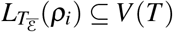, it suffices to show that 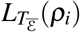 does not contain vertices of *V*(*T*) \ 𝔾. Note, 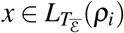 with *x* ∉ 𝔾 is only possible if all edges (*x*,*y*) are removed.

Let *x* ∈ *V* with *t*(*x*) = △ such that all edges (*x*,*y*) are removed. Thus, all such edges (*x*,*y*) are contained in ε. Therefore, every edge of the form (*x,y*) is a transfer edge; a contradiction to **(O2)**.

We will show in Prop. 1 that **(O1)**, **(O2)**, and **(O3)** together imply two important properties of event labeled species trees, **(Σ1)** and **(Σ2)**, which play a crucial role for the results reported here.

**(Σ1)** If *t*(*x*) = •, then there are distinct children *v*, *w* of *x* in *T* such that 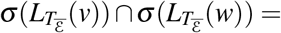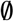.
**(Σ2)** If (*v*,*w*) ∈ ε, then 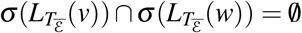.

Intuitively, **(Σ1)** is true because within a component 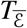 no genetic material is exchanged between non-comparable nodes. Thus, a gene separated in a speciation event necessarily ends up in distinct species in the absence of horizontal transfer. It is important to note that we do not require the converse: 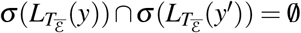 does **not** imply 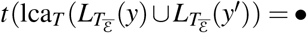 that is, the last common ancestor of two sets of genes from different species is not necessarily a speciation vertex.

Now consider a transfer edge (*v*, *w*) ∈ ε, i.e., *t*(*v*) = △. Then 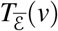 and 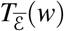 are subtrees of distinct connected components of 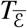. Since HGT amounts to the transfer of genetic material*across* distinct species, the genes *v* and *w* must be contained in distinct species *X* and *Y*, respectively. Since no genetic material is transferred between contemporary species *X*′ and *Y*′ in 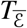, where *X*′ and *Y*′ is a descendant of *X* and *Y*, respectively we derive **(Σ1)**.

#### Proposition 1.

*Conditions **(O1)** - **(O3)** imply **(Σ1)** and **(Σ2**)*.

*Proof*. Since **(O2)** is satisfied we can apply Lemma 1 and conclude that neither 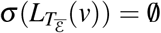 nor 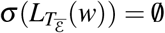. Let *x* ∈ *V*(*T*) with *t*(*x*) = •. By Condition **(O1)** *x* has (at least two) children. Moreover, **(O3)** implies that there are (at least) two children *v* and *w* in *T* that are contained in distinct species *V* and *W* that are incomparable in *S*. Note, the edges (*x*, *v*) and (*x*,*w*) remain in 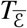, since only transfer edges are removed. Since no transfer is contained in 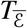, the genetic material *v* and *w* of *V* and *W*, respectively, is always vertically transmitted. Therefore, for any leaf 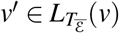 we have **σ**(*v*′) ⪯_*S*_ *V* and for any leaf 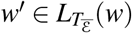 we have **σ**(*w*′) ⪯_*S*_ *W* in *S*. Assume now for contradiction, that 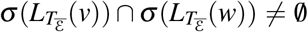. Let 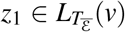 and 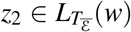 with **σ**(*z*_1_) = **σ**(*z*_2_) = *Z*. Since *Z* ⪯_*S*_ *V*,*W* and *S* is a tree, the species *V* and *W* must be comparable in *S*; a contradiction to **(O3)**. Hence, Condition **(Σ1)** is satisfied.

To see **(Σ2)**, note that since **(O2)** is satisfied we can apply Lemma 1 and conclude that neither 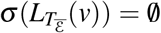 nor 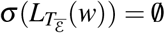. Let (*v*, *w*) ∈ ε. By **(O3)** the species containing *V* and *W* are incomparable in *S*. Now we can argue along the same lines as in the proof for **(Σ2)** to conclude that 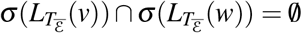.

From here on we simplify the notation a bit and write 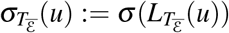. We are aware of the fact that condition **(O3)** cannot be checked directly for a given event-labeled gene tree. In contrast, **(Σ1)** and **(Σ2)** are easily determined. Hence, in the remainder of this paper we consider the more general case, that is, gene trees that satisfy **(O1)**, **(O2)**, **(Σ1)**, and **(Σ2)**.

## 4 DTL-scenario and Time-Consistent Reconciliation Maps

In case that the event-labeling of *T* is unknown, but the gene tree *T* and a species tree *S* are given, the authors in [40, 5] provide an axiom set, called DTL-scenario, to reconcile *T* with *S*. This reconciliation is then used to infer the event-labeling *t* of *T*. Instead of defining a DTL-scenario as octuple [40, 5], we use here the notation established above:

#### Definition 1

(DTL-scenario). *For a given gene tree* (*T*; *t*, **σ**) *on* 𝔾 *and a species tree S on* 𝕊 *the map γ*: *V*(*T*) → *V*(*S*) *maps the gene tree into the species tree such that*

I. *For each leaf x* ∈ 𝔾, *γ*(*u*) = **σ**(*u*).
II. *If u* ∈ *V*(*T*) \ 𝔾 *with children v*, *w*, *then*

a. *γ*(*u*) *is not a proper descendant of γ*(*v*) or *γ*(*w*), *and*
b. *at least one of γ*(*v*) *or γ*(*w*) *is a descendant of γ*(*u*).
III. (*u*, *v*) *is a transfer edge if and only if γ*(*u*) and *γ*(*v*) are incomparable.
IV. *If u* ∈ *V*(*T*) \ 𝔾 *with children v*, *w*, *then*

a. *t* (*u*) = △ *if and only if either* (*u*, *v*) *or* (*u*, *w*) *is a transfer-edge*,
b. *If t*(*u*) = •, *then γ*(*u*) = lca_*S*_(*γ*(*v*), *γ*(*w*)) and *γ*(*v*), *γ*(*w*) are incomparable,
c. *If t*(*u*) = ◻, *then γ*(*u*) ⪰ lca_*S*_ (*γ*(*v*), *γ*(*w*)).

DTL-scenarios are explicitly defined for fully resolved binary gene and species trees. Indeed, Fig. 1 (right) shows a valid reconciliation between a gene tree *T* and a species tree *S* that is not consistent with DTL-scenario. To see this, let us call the duplication vertex *v*. The vertex *v* and the leaf *a* are both children of the speciation vertex *ρ*_*T*_. Condition (IVb) implies that *a* and *v* must be incomparable. However, this is not possible since *γ*(*v*) ⪰_*S*_ lca_*S*_(*B*, *C*) (Cond. (IVc)) and *γ*(*a*) = *A* (Cond. (I)) and therefore, *γ*(*v*) ⪰_*S*_ lca_*S*_(*B*,*C*) = lca_*S*_(*A*,*B*,*C*) ≻_*S*_ *γ*(*a*).

**Figure 2:**
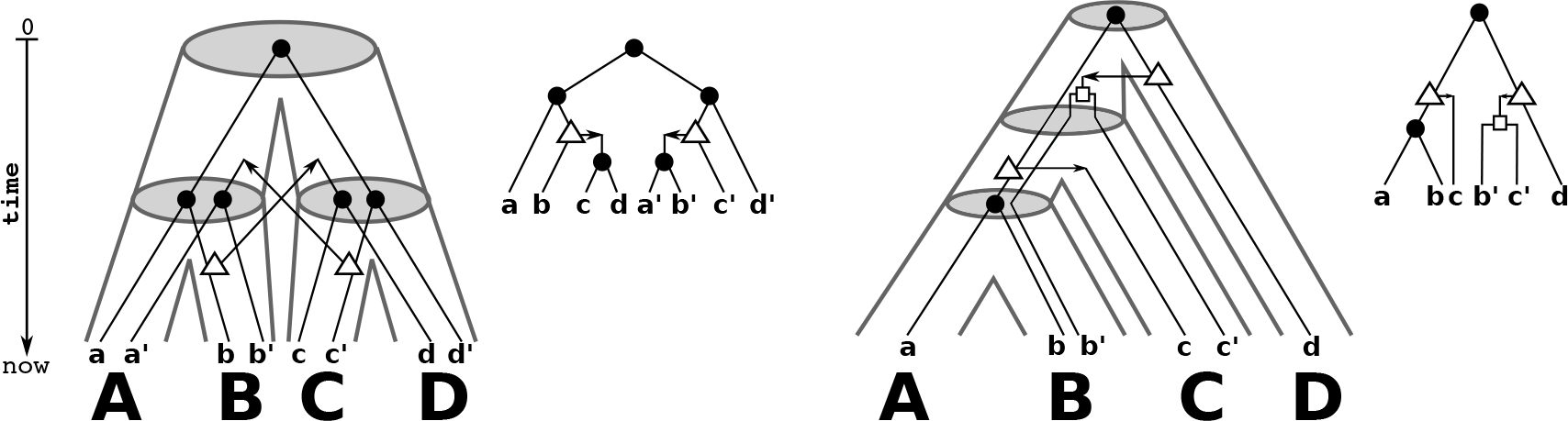
Shown are two (tube-like) species trees with reconciled gene trees. The reconciliation map *μ* for *T* and *S* is given implicitly by drawing the gene tree (upper right to the respective species tree) within the species tree. In the left example, the map *μ*, is unique. However, *μ* is not time-consistent and thus, there is no time consistent reconciliation for *T* and *S*. In the example on the right hand side, *μ*, is time-consistent.

The problem of reconciliations between gene trees and species tree is formalized in terms of so-called DTL-scenarios in the literature [40, 5]. This framework, however, usually assumes that the event labels *t* on *T* are unknown, while a species tree *S* is given. The “usual” DTL axioms, furthermore, explicitly refer to binary, fully resolved gene and species trees. We therefore use a different axiom set here that is a natural generalization of the framework introduced in [25] for the HGT-free case:

### Definition 2.

*Let T* = (*V*, *E*) *and S* = (*W*, *F*) *be phylogenetic trees on* 𝔾 and 𝕊, *resp*., **σ**: 𝔾 → 𝕊 *the assignment of genes to species and t*: *V* ⋃ *E* → {•, ◻, △, ⨀}⋃{0,1} *an event labeling on T*.

*A map μ*: *V* → *W* ⋃ *F is a reconciliation map if for all v* ∈ *V it holds that*:

**(M1)** Leaf Constraint. *If t*(*v*) = ⨀, then *μ*(*v*) = **σ**(*v*).
**(M2)** Event Constraint.

i. *If t*(*v*) = •, then 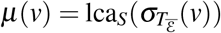.
ii. *If t*(*v*) ∈ {◻, △}, *then μ*(*v*) ∈ *F*.
iii. *If t*(*v*) = △ and (*v*, *w*) ∈ ε, *then* (*v*) *and* (*w*) *are incomparable in S*.
**(M3)** Ancestor Constraint. *Suppose v*, *w* ∈ *V with* 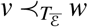.

i. *If t*(*v*), *t*(*w*) ∈ {◻, △}, *then μ*(*v*) ⪯_*S*_ *μ*(*w*),
ii. *otherwise, i.e., at least one of t*(*v*) *and t*(*w*) *is a speciation* •, *μ*(*v*) ≺_*S*_ *μ*(*w*).

*We say that S is a species tree for* (*T*; *t*, **σ**) *if a reconciliation map μ*: *V* → *W* ⋃ *F exists*.

For the special case that gene and species trees are binary, Definition 2 is equivalent to the definition of a DTL-scenario, which is summarized in the following

#### Theorem 1.

*For a binary gene tree* (*T*; *t*, **σ**) *and a binary species tree S there is a DTL-scenario if and only if there is a reconciliation for* (*T*; *t*, **σ**) *and S*.

The proof of Theorem 1 is a straightforward but tedious case-by-case analysis. In order to keep this section readable, we relegate the proof of Theorem 1 to Section 6. Figure 1 shows an example of a biologically plausible reconciliation of non-binary trees that is valid w.r.t. Definition 2 but does not satisfy the conditions of a DTL-scenario.

Condition **(M1)** ensures that each leaf of *T*, i.e., an extant gene in 𝔾, is mapped to the species in which it resides. Conditions **(M2.i)** and **(M2.ii)** ensure that each inner vertex of *T* is either mapped to a vertex or an edge in *S* such that a vertex of *T* is mapped to an interior vertex of *S* if and only if it is a speciation vertex. Condition **(M2.i)** might seem overly restrictive, an issue to which we will return below. Condition **(M2.iii)** satisfies condition **(O3)** and maps the vertices of a transfer edge in a way that they are incomparable in the species tree, since a HGT occurs between distinct (co-existing) species. It becomes void in the absence of HGT; thus Definition 2 reduces to the definition of reconciliation maps given in [25] for the HGT-free case. Importantly, condition **(M3)** refers only to the connected components of 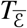 since comparability w.r.t. 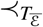 implies that the path between *x* and *y* in *T* does not contain transfer edges. It ensures that the ancestor order ⪯_*T*_ of *T* is preserved along all paths that do not contain transfer edges.

We will make use of the following bound that effectively restricts how close to the leafs the image of a vertex in the gene tree can be located.

### Lemma 2.

*If μ*: (*T*; *t*, **σ**) → *S satisfies **(M1)** and **(M3)**, then* 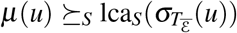 *for any u* ∈ *V*(*T*).

*Proof*. If *u* is a leaf, then by Condition **(M1)** *μ*(*u*) = **σ**(*u*) and we are done. Thus, let *u* be an interior vertex. By Condition **(M3)**, *z* ⪯_*S*_ *μ*(*u*) for all 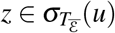. Hence, if 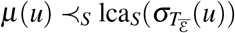 or if *μ*(*u*) and 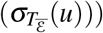 are incomparable in *S*, then there is a 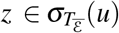) such that *z* and *μ*(*u*) are incomparable; contradicting **(M3)**.

Condition **(M2.i)** implies in particular the weaker property “(M2.i’) if *t*(*v*) = • then *μ*(*v*) ∈ *W*”. In the light of Lemma 2, 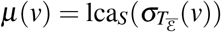 is the lowest possible choice for the image of a speciation vertex. Clearly, this restricts the possibly exponentially many reconciliation maps for which 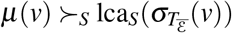 for a speciation vertices *v* to only those that satisfy **(M2.i)**. However, the latter is justified by the observation that if *v* is a speciation vertex with children *u*, *w*, then there is only one unique piece of information given by the gene tree to place *μ*(*v*), that is, the unique vertex *x* in *S* with children *y*, *z* such that 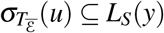 and 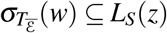. The latter arguments easily generalizes to the case that *v* has more than two children in *T*. Moreover, any *observable* speciation node *v*′ ≻_*T*_ *v* closer to the root than *v* must be mapped to a node ancestral to *μ*(*v*) due to **(M3.ii)**. Therefore, we require 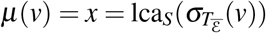 here.

If *S* is a species tree for the gene tree (*T*, *t*, **σ**) then there is no freedom in the construction of a reconciliation map *μ* on the set {*x* ∈ *V*(*T*) | *t*(*x*) ∈ {•, ⨀}}. The duplication and HGT vertices of *T*, however, can be placed differently. As a consequence there is a possibly exponentially large set of reconciliation maps from (*T*, *t*, **σ**) to *S*.

From a biological point of view, however, the notion of reconciliation used so far is too weak. In the absence of HGT, subtrees evolve independently and hence, the linear order of points along each path from root to leaf is consistent with a global time axis. This is no longer true in the presence of HGT events, because HGT events imply additional time-consistency conditions. These stem from the fact that the appearance of the HGT copy in a distant subtree of *S* is concurrent with the HGT event. To investigate this issue in detail, we introduce time maps and the notion of time-consistency, see Figures 2-4 for illustrative examples.

**Definition 3** (Time Map). *The map* τ_*T*_: *V*(*T*) → ℝ *is a time map for the rooted tree T if x* ≺_*T*_ y *implies* τ_*T*_(*x*) > τ_*T*_(*y*) *for all x*,*y* ∈ *V*(*T*).

**Definition 4.** *A reconciliation map μ from* (*T*; *t*, **σ**) *to S is time-consistent if there are time maps* τ_*T*_ for *T* and τ_*S*_ *for S for all u* ∈ *V*(*T*) *satisfying the following conditions*:

**(C1)** *If t*(*u*) ∈ {•, ⨀}, *then* τ_*T*_ (*u*) = τ_*S*_(*μ*(*u*)).
**(C2)** *If t*(*u*) ∈ {◻, △} *and, thus μ*(*u*) = (*x*, *y*) ∈ *E*(*S*), then τ_*S*_(y) > τ_*T*_(*u*) > τ_*S*_(*x*).

Condition **(C1)** is used to identify the time-points of speciation vertices and leaves *u* in the gene tree with the time-points of their respective images *μ*(*u*) in the species trees. In particular, all genes *u* that reside in the same species must be assigned the same time point τ_*T*_(*u*) = τ_*S*_(**σ**(*u*)). Analogously, all speciation vertices in *T* that are mapped to the same speciation in *S* are assigned matching time stamps, i.e., if *t*(*u*) = *t*(*v*) = • and *μ*(*u*) = *μ*(*v*) then τ_*T*_(*u*) = τ_*T*_(*v*) = τ_*S*_(*μ*(*u*)).

**Figure 3:**
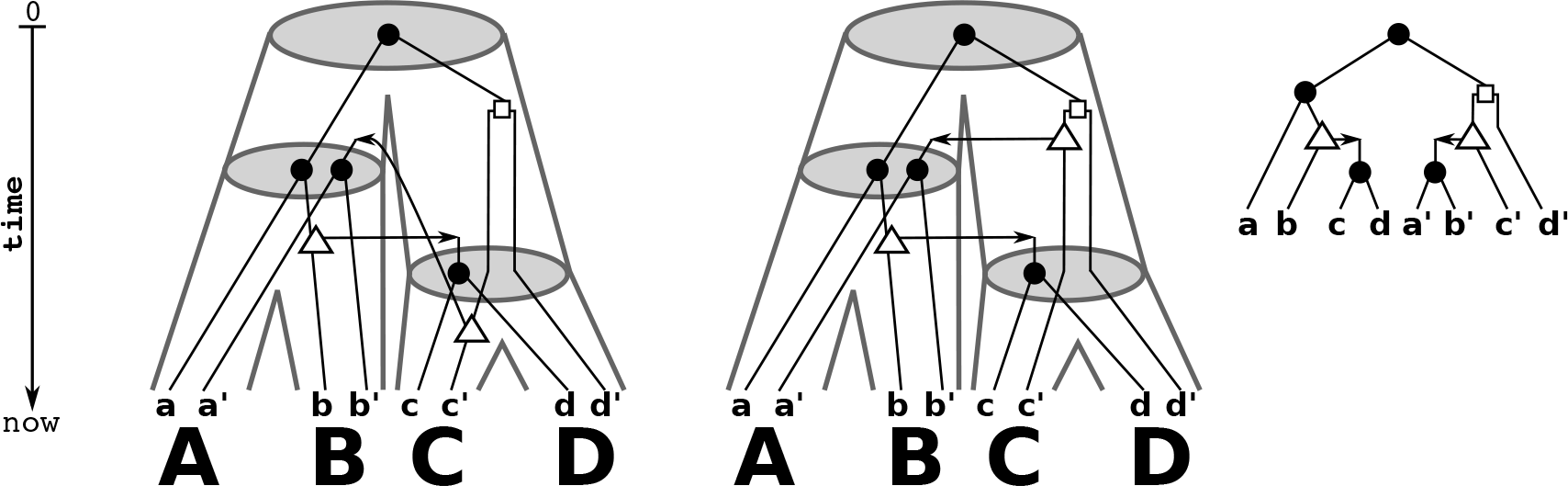
Shown are a gene tree (*T*; *t*, **σ**) (right) and two identical (tube-like) species trees *S* (left and middle). There are two possible reconciliation maps for *T* and *S* that are given implicitly by drawing *T* within the species tree *S*. These two reconciliation maps differ only in the choice of placing the HGT-event either on the edge (lca_*S*_(*C*,*D*),*C*) or on the edge (lca_*S*_({*A*,*B*,*C*,*D*}), lcas(*C*,*D*)). In the first case, it is easy to see that *μ*, would not be time-consistent, i.e., there are no time maps τ_*T*_ and τ_*S*_ that satisfy **(C1)** and **(C2)**. The reconciliation map *μ*, shown in the middle is time-consistent.

**Figure 4:**
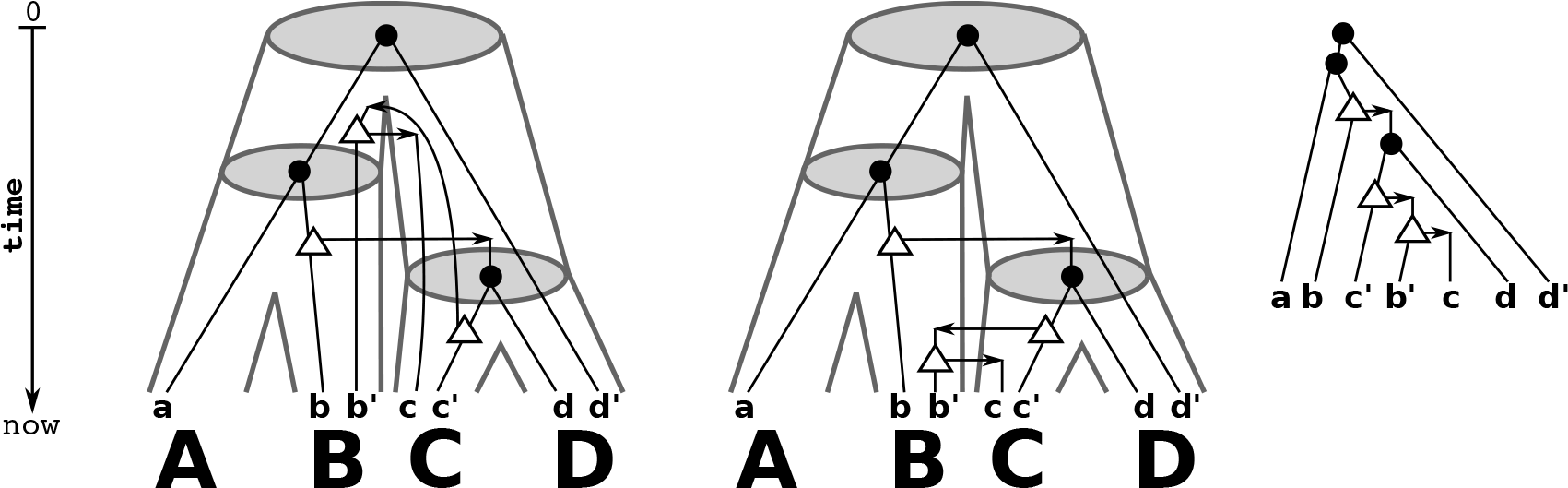
Shown are a gene tree (*T*; *t*, **σ**) (right) and two identical (tube-like) species trees *S* (left and middle). There are two possible reconciliation maps for *T* and *S* that are given implicitly by drawing *T* within the species tree *S*. The left reconciliation maps each gene tree vertex as high as possible into the species tree. However, in this case only the middle reconciliation map is time-consistent.

To understand the intuition behind **(C2)** consider a duplication or HGT vertex *u*. By construction of *μ* it is mapped to an edge of *S*, i.e., *μ*(*u*) = (*x*,*y*) in *S*. The time point of *u* must thus lie between time points of *x* and *y*. Now suppose (*u*, *v*) ∈ ε is a transfer edge. By construction, *u* signifies the transfer event itself. The node *v*, however, refers to the next (visible) event in the gene tree. Thus τ_*T*_ (*u*) < τ_*T*_(*v*). In particular, τ_*T*_(*v*) must not be misinterpreted as the time of introducing the HGT-duplicate into the new lineage. While this time of course exists (and in our model coincides with the timing of the transfer event) it is not marked by a visible event in the new lineage, and hence there is no corresponding node in the gene tree *T*.

W.l.o.g. we fix the time axis so that τ_*T*_(***ρ***_*T*_) = 0 and τ_*S*_(***ρ***_*S*_) = −1. Thus, τ_*s*_(***ρ***_*S*_) < τ_*T*_(***ρ***_*T*_) < τ_*T*_(*u*) for all *u* ∈ *V*(*T)*) \ {***ρ***_*T*_}.

Clearly, a necessary condition to have biologically feasible gene trees is the existence of a reconciliation map *μ*. However, not all reconciliation maps are time-consistent, see Fig. 2.

### Definition 5.

*An event-labeled gene tree* (*T*; *t*, **σ**) *is* biologically feasible *if there exists a time-consistent reconciliation map from* (*T*; *t*, **σ**) *to some species tree S*.

As a main result of this contribution, we provide simple conditions that characterize (the existence of) time-consistent reconciliation maps and thus, provides a first step towards the characterization of biologically feasible gene trees.

### Theorem 2.

*Let μ be a reconciliation map from* (*T; t, σ*) *to S. There is a time-consistent reconciliation map from* (*T; t, σ*) *to S if and only if there are two time-maps τ*_*T*_ *and τ*_*S*_ *for T and S, respectively, such that the following conditions are satisfiedfor all x* ∈ *V*(*S*):

**(D1)** *If μ*(*u*) = *x*, *for some u* ∈ *V*(*T)*), *then* τ_*T*_(*u*) = τ_*S*_(*x*).
**(D2)** *If* 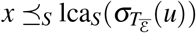 *for some u* ∈ *V*(*T*) *with t*(*u*) ∈ {◻, △}, then τ_*s*_(*x*) > τ_*T*_(*u*).
**(D3)** *If* 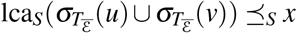 *for some* (*u*, *v*) ∈ ε, *then* τ_*T*_(*u*) > τ_*s*_(*x*).

*Proof*. In what follows, *x* and *u* denote vertices in *S* and *T*, respectively.

Assume that there is a time-consistent reconciliation map *μ* from (*T*; *t*, **σ**) to *S*, and thus two time-maps τ_*s*_ and τ_*T*_ for *S* and *T*, respectively, that satisfy **(C1)** and **(C2)**.

To see **(D1)**, observe that if *μ*(*u*) = *x* ∈ *V*(*S*), then **(M1)** and **(M2)** imply that *t*(*u*) ∈ {•, ⨀}. Now apply **(C1)**.

To show **(D2)**, assume that *t*(*u*) ∈ {◻, △} and 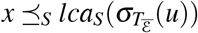. By Condition **(M2)** it holds that *μ*(*u*) = (*y*, *z*) ∈ *E*(*S*). Together with Lemma 2 we obtain that *x* 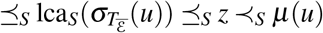. By the properties of τ_*S*_ we have

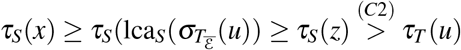

To see **(D3)**, assume that (*u*, *v*) ∈ ε and 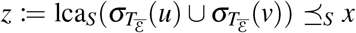. Since *t*(*u*) = △ and by **(M2ii)**, we have *μ*(*u*) = (*y*,*y*′) ∈ *E*(*S*). Thus, *μ*(*u*) ≺_s_ *y*. By **(M2iii)** *μ*(*u*) and *μ*(v) are incomparable and therefore, we have either *μ* (v) ≺_*s*_ *y* or *μ*(*v*) and *y* are incomparable. In either case we see that *y* ⪯_*S*_ *z*, since Lemma 2 implies that 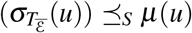 and 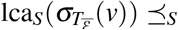 *μ* (*v*). In summary, *μ*(*u*) ≺_*S*_ *y* ⪯_*S*_ *z* ⪯_*S*_ *x*. Therefore,

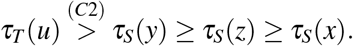

Hence, conditions **(D1)**-**(D3)** are satisfied.

To prove the converse, assume that there exists a reconciliation map *μ* that satisfies **(D1)**-**(D3)** for some time-maps τ_*T*_ and τ_*s*_. In the following we will make use of τ_*s*_ and τ_*T*_ to construct a time-consistent reconciliation map *μ*′.

First we define “anchor points” by *μ*′(*v*) = *μ*(*v*) for all *v* ∈ *V*(*T*) with *t*(*v*) ∈ {•, ⨀}. Condition **(D1)** implies τ_*T*_(*v*) = τ_*s*_(*μ*(*v*)) for these vertices, and therefore *μ*′ satisfies **(C1)**.

The next step will be to show that for each vertex *u* ∈ *V*(*T*) with *t*(*u*) ∈ {◻, △} there is a unique edge (*x*,*y*) along the path from 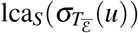 to ***ρ***_*S*_ with τ_*s*_(*x*) < τ_*T*_(*u*) < τ_*S*_(*y*). We set *μ*′(*u*) = (*x*,*y*) for these points. In the final step we will show that *μ*′ is a valid reconciliation map.

Consider the unique path 𝒫_*u*_ from 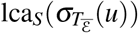 to ***ρ***_*S*_. By construction, τ_*s*_(***ρ***_*S*_) < τ_*T*_(***ρ***_*T*_) ≤ τ_*T*_(*u*) and by Condition **(D2)** it we have 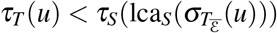. Since τ_*s*_ is a time map for *S*, every edge (*x*,*y*) ∈ *E*(*S*) satisfies τ_*s*_(*x*) < τ_*S*_(*y*). Therefore, there is a unique edge (*x*_*u*_,*y*_*u*_) ∈ *E*(*S*) along 𝒫_*u*_ such that either τ_s_(*x*_*u*_) < τ_T_(*u*) < τ_S_(*y*_*u*_), τ_S_(*x*_*u*_) = τ_T_(*u*) < τ_S_(*y*_*u*_), or τ_S_(*x*_*u*_) < τ_T_(*u*) = τ_*S*_(*y*_*u*_). The addition of a sufficiently small perturbation ε_*u*_ to τ_*T*_ (*u*) does not violate the conditions for τ_*T*_ being a time-map for *T*. Clearly ε_*u*_ can be chosen to break the equalities in the latter two cases in such a way that τ_*s*_(*x*_*u*_) < τ_*T*_(*u*) < τ_*S*_(*y*_*u*_) for each vertex *u* ∈ *V*(*T*) with *t*(*u*) ∈ {◻, △}. We then continue with the perturbed version of τ_*T*_ and set *μ*′(*u*) = (*x*_*u*_,*y*_*u*_). By construction, *μ*′ satisfies **(C2)**.

It remains to show that *μ*′ is a valid reconciliation map from 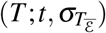 to *S*. Again, let 𝒫_*u*_ denote the unique path from 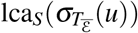 to ***ρ***_*S*_ for any *u* ∈ *V*(*T*).

By construction, Conditions **(M1)**, **(M2i)**, **(M2ii)** are satisfied. To check condition **(M2iii)**, assume (*u*, *v*) ∈ ε. The original map *μ* is a valid reconciliation map, and thus, Lemma 2 implies that 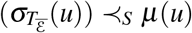 and 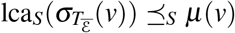. Since *μ*(*u*) and *μ*(*v*) are incomparable in *S* and 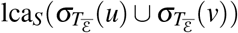 lies on both paths 𝒫_*u*_ and 𝒫_*v*_ we have *μ*(*u*), 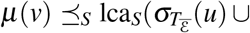 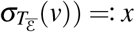. In particular, 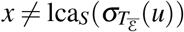 and 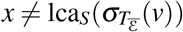.

Conditions **(D1)** and **(D2)** imply that 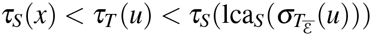 and 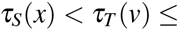 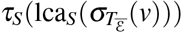. By construction of *μ*′, the vertex *u* is mapped to a unique edge *e*_*u*_ = (*x*_*u*_,*y*_*u*_) and *v* is mapped either to 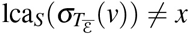 or to the unique edge *e*_*v*_ = (*x*_*v*_,*y*_*v*_), respectively. In particular, *μ*′(*u*) lies on the path 𝒫′ from *x* to 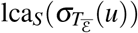 and *μ*′(*v*) lies one the path 𝒫″ from *x* to 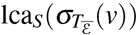. The paths 𝒫′ and 𝒫″ are edge-disjoint and have *x* as their only common vertex. Hence, *μ*′(*u*) and *μ*′(*v*) are incomparable in *S*, and **(M2iii)** is satisfied.

In order to show **(M3)**, assume that 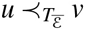. Since 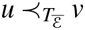, we have 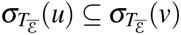. Hence, 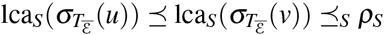. In other words, 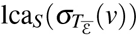 lies on the path 𝒫_*u*_ and thus, 𝒫_*v*_ is a subpath of 𝒫_*u*_. By construction of *μ*′, both *μ*′(*u*) and *μ*′(*v*) are comparable in *S*. Moreover, since τ_*T*_(*u*) > τ_*T*_(*v*) and by construction of *μ*′, it immediately follows that *μ*′(*u*) ⪯_*S*_ *μ*′(*v*).

Its now an easy task to verify that **(M3)** is fulfilled by considering the distinct event-labels in **(M3i)** and **(M3ii)**, which we leave to the reader.

Interestingly, the existence of a time-consistent reconciliation map from a gene tree *T* to a species tree *S* can be characterized in terms of a time map defined on *T*, only.

### Theorem 3.

*Let μ be a reconciliation map from* (*T*; *t*, **σ**) *to S. There is a time-consistent reconciliation map* (*T*; *t*, **σ**) *to S if and only if there is a time map* τ_*T*_ *such that for all u*, *v*, *w* ∈ *V*(*T*):

**(T1)** *If t*(*u*) = *t*(*v*) ∈ {•, ⨀} *then*

a. *If μ*(*u*) = *μ*(*v*), *then* τ_*T*_ (*u*) = τ_*T*_(*v*).
b. *If μ*(*u*) ≺_*S*_ *μ*(*u*), *then* τ_*T*_(*u*) > τ_*T*_(*v*).
**(T2)** *If t*(*u*) ∈ {•, ⨀}, *t*(*v*) ∈ {◻, △} *and* 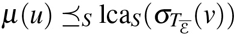, *then* τ_*T*_(*u*) > τ_*T*_(*v*).
**(T3)** *If* (*u*, *v*) ∈ ε and 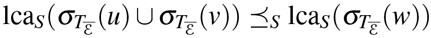 *for some w* ∈ *V*(*T*), then τ_*T*_(*u*) > τ_*T*_(*w*)

*Proof*. Suppose that *μ* is a time-consistent reconciliation map from (*T*; *t*, **σ**) to *S*. By Definition 4 and Theorem 2, there are two time maps τ_*T*_ and τ_*s*_ that satisfy **(D1)**-**(D3)**. We first show that τ_*T*_ also satisfies **(T1)**-**(T3)**, for all *u*, *v* ∈ *V*(*T*). Condition **(T1a)** is trivially implied by **(D1)**. Let *t*(*u*),*t*(*v*) ∈ {•, ⨀}, and *μ*(*u*) ≺_*S*_ *μ*(*v*). Since τ_*T*_ and τ_*s*_ are time maps, we may conclude that

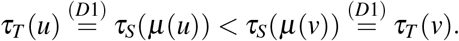

Hence, **(T1b)** is satisfied.

Now, assume that *t*(*u*) ∈ {•, ⨀}, *t*(*v*) ∈ {◻, △} and 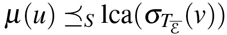. By the properties of τ_S_, we have:

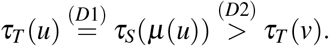

Hence, **(T2)** is fulfilled.

Finally, assume that (*u*, *v*) ∈ ε, and 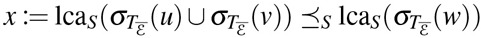 for some *w* ∈ *V*(*T*). Lemma 2 implies that 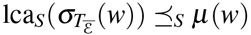 and we obtain

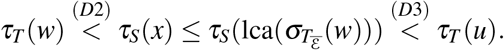

Hence, **(T3)** is fulfilled.

To see the converse, assume that there exists a reconciliation map *μ* that satisfies **(T1)**-**(T3)** for some time map τ_*T*_. In the following we construct a time map τ_*s*_ for *S* that satisfies **(D1)**-**(D3)**. To this end, we first set

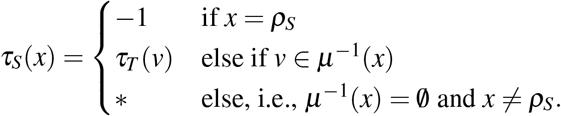

We use the symbol ∗ to denote the fact that so far no value has been assigned to τ_*s*_(*x*). Note, by **(M2i)** and **(T1a)** the value τ_*S*_(*x*) is uniquely determined and thus, by construction, **(D1)** is satisfied. Moreover, if *x*, *y* ∈ *V* (*S*) have non-empty preimages w.r.t. *μ* and *x* ≺_*S*_ *y*, then we can use the fact that τ_*T*_ is a time map for *T* together with condition **(T1)** to conclude that τ_*S*_(*x*) > τ_*S*_(*y*).

If *x* ∈ *V*(*S*) with *a* ∈ *μ*^−1^(*x*), then **(T2)** implies **(D2)** (by **(D1)** and setting *u* = *a* in **(T2)**) and **(T3)** implies **(D3)** (by **(D1)** and setting *w* = *a* in **(T3)**). Thus, **(D2)** and **(D3)** is satisfied for all *x* ∈ *V*(*S*) with 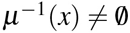.

Using our choices τ_*S*_(***ρ***_*T*_) = 0 and τ_S_(***ρ***_*S*_) = −1 for the augmented root of *S*, we must have 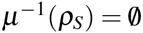. Thus, 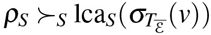 for any *v* ∈ *V*(*T*). Hence, **(D2)** is trivially satisfied for ***ρ***_*S*_. Moreover, τ_*T*_(***ρ***_*T*_) = 0 implies τ_*T*_(*u*) > τ_*s*_(***ρ***_*S*_) for any *u* ∈ *V*(*T*). Hence, **(D3)** is always satisfied for ***ρ***_*S*_.

In summary, Conditions **(D1)**-**(D3)** are met for any vertex *x* ∈ *V*(*S*) that up to this point has been assigned a value, i.e., τ_*S*_(*x*) ≠ ∗.

We will now assign to all vertices *x* ∈ *V*(*S*) with 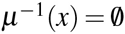 a value τ_*S*_(*x*) in a stepwise manner. To this end, we give upper and lower bounds for the possible values that can be assigned to τ_*S*_(*x*). Let *x* ∈ *V*(*S*) with τ_*S*_(*x*) = ∗. Set

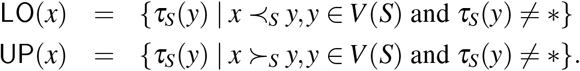

We note that 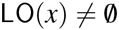 and 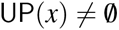 because the root and the leaves of *S* already have been assigned a value τ_*s*_ in the initial step. In order to construct a valid time map τ_*s*_ we must ensure max(LO(*x*)) < τ_*s*_(*x*) < min(UP(*x*)).

Moreover, we strengthen the bounds as follows. Put

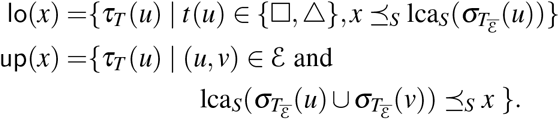

Observe that max(lo(*x*)) < min(up(*x*)), since otherwise there are vertices *u*,*w* ∈ *V*(*T*) with τ_*T*_(*w*) ∈ lo(*x*) and τ_*T*_(*u*) ∈ up(*x*) and τ_*T*_(*w*) ≥ τ_*T*_(*u*). However, this implies that 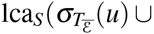 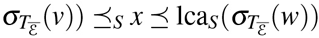 a contradiction to **(T3)**.

Since **(D2)** is satisfied for all vertices y that obtained a value τ_*S*_(*y*) ≠ ∗, we have max(lo(*x*)) < min(UP(*x*)). Likewise because of **(D3)**, it holds that max(LO(*x*)) < min(up(*x*)). Thus we set τ_*S*_(*x*) to an arbitrary value such that

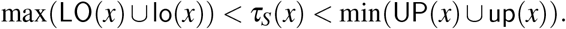

By construction, **(D1)**, **(D2)**, and **(D3)** are satisfied for all vertices in *V*(*S*) that have already obtained a time value distinct from −. Moreover, for all such vertices with *x* ≺_*T*_ *y* we have τ_*S*_(*X*) > τ_*S*_(*y*). In each step we chose a vertex *x* with τ_*S*_(*x*) = ∗ that obtains then a real-valued time stamp. Hence, in each step the number of vertices that have value ∗ is reduced by one. Therefore, repeating the latter procedure will eventually assign to all vertices a real-valued time stamp such that, in particular, τ_*s*_ satisfies **(D1)**, **(D2)**, and **(D3)** and thus is indeed a time map for *S*.

From the algorithmic point of view it is desirable to design methods that allow to check whether a reconciliation map is time-consistent. Moreover, given a gene tree *T* and species tree *S* we wish to decide whether there exists a time-consistent reconciliation map *μ*, and if so, we should be able to construct *μ*.

To this end, observe that any constraints given by Definition 3, Theorem 2 **(D2)**-**(D3)**, and Definition 4 **(C2)** can be expressed as a total order on *V*(*S*) ⋃ *V*(*T*), while the constraints **(C1)** and **(D1)** together suggest that we can treat the preimage of any vertex in the species tree as a “single vertex”. In fact we can create an auxiliary graph in order to answer questions that are concerned with time-consistent reconciliation map *S*.

### Definition 6.

*Let μ be a reconciliation map from* (*T*; *t*, σ) *to S. The auxiliary graph A is defined as a directed graph with a vertex set V* (*A*) = *V*(*S*) ⋃ *V*(*T*) *and an edge-set E*(*A*) *that is constructed as follows*

**(A1)** For each (*u*, *v*) ∈ *E*(*T*) we have (*u*′, *v*′) ∈ *E*(*A*), where

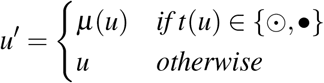

*and*

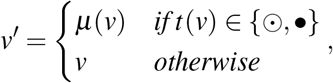
**(A2)** *For each* (*x*,*y*) ∈ *E*(*S*) *we have* (*x*,*y*) ∈ *E*(*A*).
**(A3)** *For each u* ∈ *V*(*T*) with *t*(*u*) ∈ {◻, △} *we have* 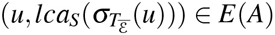.
**(A4)** *For each* (*u*, *v*) ∈ ε *we have* 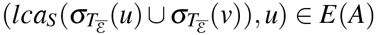
**(A5)** *For each u* ∈ *V*(*T*) *with t*(*u*) ∈ {△, ◻} and *μ*(*u*) = (*x*, *y*) ∈ *E*(*S*) *we have* (*x*, *u*) ∈ *E*(*A*) an*d* (*u*,*y*) ∈ *E*(*A*).

*We define A*_1_ *and A*_*2*_ *as the subgraphs of A that contain only the edges defined by **(A1)**, **(A2)**, **(A5)** and **(A1)**, **(A2)**, **(A3)**, **(A4)**, respectively.*

We note that the edge sets defined by conditions **(A1)** through **(A5)** are not necessarily disjoint. The mapping of vertices in *T* to edges in *S* is considered only in condition **(A5)**. The following two theorems are the key results of this contribution.

### Theorem 4.

*Let μ be a reconciliation map from* (*T*; *t*, **σ**) *to S. The map μ is time-consistent if and only if the auxiliary graph A_1_ is a directed acyclic graph (DAG).*

*Proof*. Assume that *μ* is time-consistent. By Theorem 2, there are two time-maps τ_*T*_ and τ_*S*_ satisfying **(C1)** and **(C2)**. Let τ = τ_*T*_ ⋃ τ_*S*_ be the map from *V*(*T*) ⋃ *V*(*S*) → ℝ. Let *A*′ be the directed graph with *V*(*A*′) = *V*(*S*) ⋃ *V*(*T*) and set for all *x*, *y* ∈ *V*(*A*′): (*x*, *y*) ∈ *E*(*A*′) if and only if τ(*x*) < τ(*y*). By construction *A*′ is a DAG since *τ* provides a topological order on *A*′ [26].
We continue to show that A′ contains all edges of *A*_1_.

To see that **(A1)** is satisfied for *E*(*A*′) let (*u*, *v*) ∈ *E*(*T*). Note, *τ*(*v*) > τ(*u*), since τ_*T*_ is a time map for *T* and by construction of τ. Hence, all edges (*u*, *v*) ∈ *E*(*T*) are also contained in *A*′, independent from the respective event-labels *t*(*u*), *t*(*v*). Moreover, if *t*(*u*) or *t*(*v*) are speciation vertices or leaves, then *(C1)* implies that τ_*S*_(*μ*(*u*)) = τ_*T*_(*u*) > τ_*T*_(*v*) or τ_*T*_(*u*) > τ_*T*_(*v*) = τ_*S*_(*μ*(*v*)). By construction of τ, all edges satisfying **(A1)** are contained in ∈ (*A*′). Since τ_*S*_ is a time map for s, all edges as in **(A2)** are contained in *E* (*A*′). Finally, **(C2)** implies that all edges satisfying **(A5)** are contained in ∈ (*A*′).

Although, *A*′ might have more edges than required by **(A1)**, **(A2)** and **(A5)**, the graph *A*_1_ is a subgraph of *A*′. Since *A*′ is a DAG, also *A*_1_ is a DAG.

For the converse assume that *A*_1_ is a directed graph with *V*(*A*_1_) = *V*(*S*) ⋃ *V*(*T*) and edge set *E*(*A*_1_) as constructed in Def. 6 **(A1)**, **(A2)** and **(A5)**. Moreover, assume that *A*_1_ is a DAG. Hence, there is is a topological order τ on *A*_1_ with τ(*x*) < τ(*y*) whenever (*x*,*y*) ∈ *E*(*A*_1_). In what follows we construct the time-maps τ_*T*_ and τ_*S*_ such that they satisfy **(C1)** and **(C2)**. Set τ_*S*_(*X*) = τ(*x*) for all *x* ∈ V (*S*). Additionally, set for all *u* ∈ *V*(*T*):

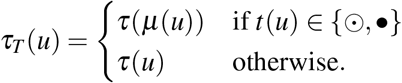

By construction it follows that **(C1)** is satisfied. Due to **(A2)**, τ_*S*_ is a valid time map for *S*. It follows from the construction and **(A1)** that τ_*T*_ is a valid time map for *T*. Assume now that *u* ∈ *V*(*T*), *t*(*u*) ∈ {◻, △}, and *μ*(*u*) = (*x*,*y*) ∈ *E*(*S*). Since τ provides a topological order we have:

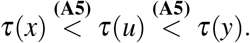

By construction, it follows that τ_*s*_(*x) < τ_*T*_(*u*) < τ_*S*_(y*) satisfying **(C2)**.

### Theorem 5.

*Assume there is a reconciliation map μ from (*T*; *t*, **σ**) to S. There is a time-consistent reconciliation map, possibly different from i, from (*T*; *t*, **σ**) to S if and only if the auxiliary graph A_*2*_ (defined on *μ*) is a DAG*.

*Proof*. Let *μ* be a reconciliation map for (*T*; *t, σ*) and *S* and *μ*′ be a time-consistent reconciliation map for (*T*; *t*, **σ**) and *S*. Let *A*_*2*_ and A′_2_ be the auxiliary graphs that satisfy Def. 6 **(A1)** - **(A4)** for *μ* and *μ*′, respectively. Since *μ*(*u*) = *μ*′(*u*) for all *u* ∈ *V*(*T*) with *t*(*u*) ∈ {⨀, •} and **(A2)** - **(A4)** don’t rely on the explicit reconciliation map, it is easy to see that *A*_*2*_ = *A*′_2_.

Now we can re-use similar arguments as in the proof of Theorem 4. Assume there is a time-consistent reconciliation map (*T*; *t*, **σ**) to *S*. By Theorem 2, there are two time-maps τ_*T*_ and τ_*s*_ satisfying **(D1)**-**(D3)**. Let *T* and *A*′ be defined as in the proof of Theorem 4.

Analogously to the proof of Theorem 4, we show that *A*^′^ contains all edges of *A*_2_. Application of **(D1)** immediately implies that all edges satisfying **(A1)** and **(A2)** are contained in *E*(*A*′). By condition **(D2)**, it yields 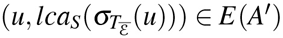 and **(D3)** implies 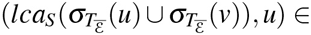 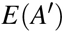. We conclude by the same arguments as before that the graph *A*_*2*_ is a DAG.

For the converse, assume we are given the directed acyclic graph *A*_2_. As before, there is is a topological order τ on *A*_2_ with τ(*x*) < τ(*y*) only if (*x,y*) ∈ *E*(*A*_2_). The time-maps τ_*T*_ and τ_*s*_ are given as in the proof of Theorem 4.

By construction, it follows that **(D1)** is satisfied. Again, by construction and the Properties **(A1)** and **(A2)**, τ_*s*_ and τ_*T*_ are valid time-maps for *S* and *T* respectively.

Assume now that *u* ∈ *V*(*T*),*t*(*u*) ∈ {◻, △}, and 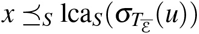 for some *x* ∈ *V*(*S*). Since there is a topological order on *V*(*A*_2_), we have

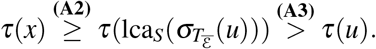

By construction, it follows that τ_*s*_(*x*) > τ_*T*_(*u*). Thus, **(D2)** is satisfied.

Finally assume that (*u*, *v*) ∈ ε and 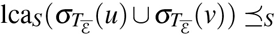 for some *x* ∈ *V*(*S*). Again, since τ provides a topological order, we have:

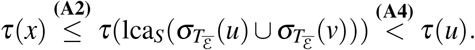

By construction, it follows that τ_*s*_(*x*) < τ_*T*_(*u*), satisfying **(D3)**.

Thus τ_*T*_ and τ_*S*_ are valid time maps satisfying **(D1)**-**(D3)**.

Naturally, Theorems 4 or 5 can be used to devise algorithms for deciding time-consistency. To this end, the efficient computation of 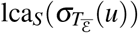 for all *u* ∈ *V*(*T*) is necessary. This can be achieved with Algorithm 2 in *O*(|*V*(*T*)|log(|*V*(*S*)|)). More precisely, we have the following statement:

#### Lemma 3.

*For a given gene tree (*T* = (*V*,*E*):*t*, **σ**) and a species tree *S* = (*W*,*F*), Algorithm 2 correctly computes 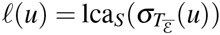 for all u* ∈ *V*(*T*) *in O*(|*V*|log(|*W*|)) *time*.

*Proof*. Let *u* ∈ *V*(*T*). In what follows, we show that 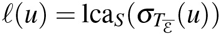. In fact, the algorithm is (almost) a depth first search through *T* that assigns the (species tree) vertex ℓ(*v*) to *u* if and only if every child *v* of *u* has obtained an assignment ℓ(*v*) (cf. Line (9) - (10)). That there are children *v* with non-empty ℓ(*v*) at some point is ensured by Line (7). That is, if *t*(*u*) = ⨀, then 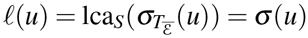. Now, assume there is an interior vertex *u* ∈ *V*(*T*), where every child *v* has been assigned a value ℓ(*v*), then

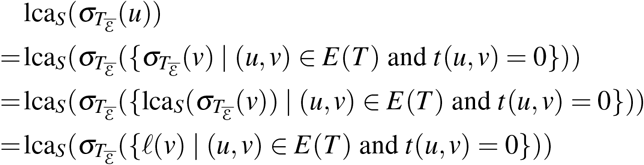

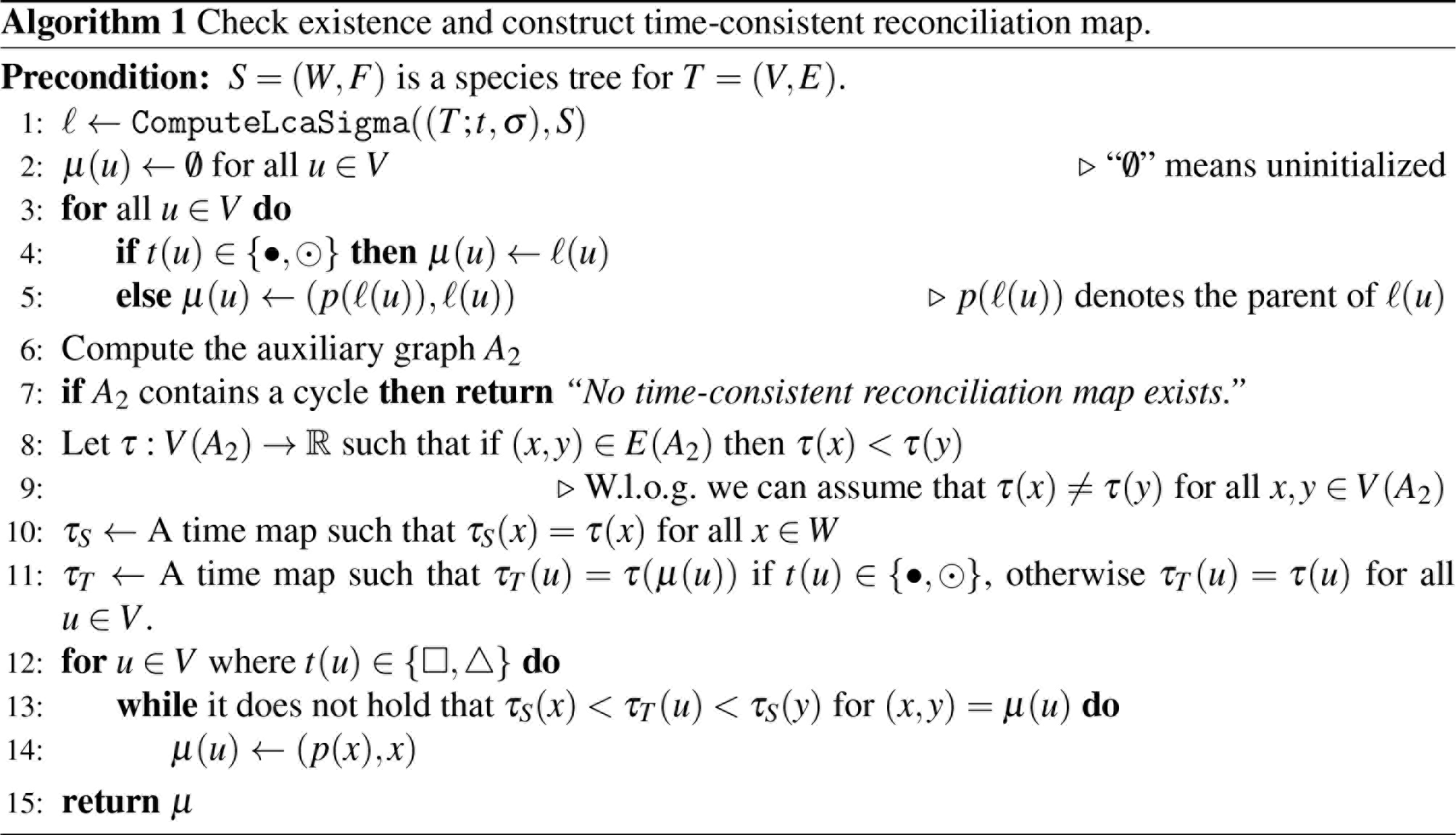

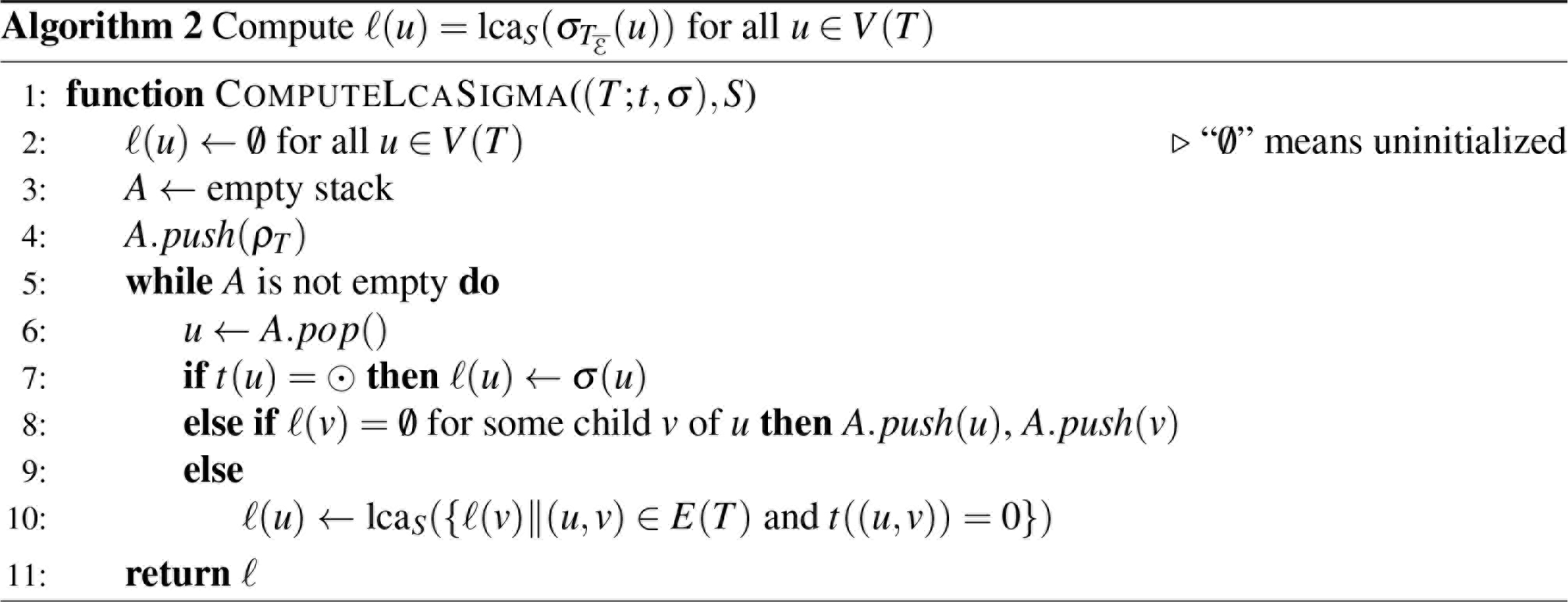

The latter is achieved by Line (10).

Since *T* is a tree and the algorithm is in effect a depth first search through *T*, the while loop runs at most *O*(*V*(*T*) + *E*(*T*)) times, and thus in *O*(*V*(*T*)) time.

The only non-constant operation within the while loop is the computation of lca_*S*_ in Line (10). Clearly lca_*S*_ of a set of vertices *C* = {*c*_1_, *c*_2_ … *c*_*k*_}, where *c*_*i*_ ∈ *V*(*S*), for all *c*_*i*_ ∈ *C* can be computed as sequence of lca_*S*_ operations taking two vertices: lca_*S*_(*c*_1_, lca_*S*_(*c*_2_,… lca_*S*_(*c*_*k−1*_, *c*_*k*_))), each taking *O*(lg(|*V*(*S*)|)) time. Note however, that since Line (10) is called exactly once for each vertex in *T*, the number of lca_*S*_ operations taking two vertices is called at most |*E*(*T*)| times through the entire algorithm. Hence, the total time complexity is *O*(|*V*(*T*)|lg(|*V*(*S*)|)).

Let *S* be a species tree for (*T*; *t*, **σ**), that is, there is a valid reconciliation between the two tree*S*. Algorithm 1 describes a method to construct a time-consistent reconciliation map for (*T*; *t*, **σ**) and *S*, if one exists, else “No time-consistent reconciliation map exists” is returned. First, an arbitrary reconciliation map *μ* that satisfies the condition of Def. 2 is computed. Second, Theorem 5 is utilized and it is checked whether the auxiliary graph *A*_2_ is not a DAG in which case no time-consistent map *μ* exists for (*T*; *t*, **σ**) and *S*. Finally, if *A*_2_ is a DAG, then we continue to adjust *μ* to become time-consistent. The latter is based on Thm. 2, see the proof of Thm. 2 and 6 for detail*S*.

#### Theorem 6.

*Let S = (W,F) be species tree for the gene tree (T = (V,E); t, **σ**). Algorithm 1 correctly determines whether there is a time-consistent reconciliation map *μ* and in the positive case, returns such a μ* in *O*(|*V*| log(|*W*|)) *time*.

*Proof*. In order to produce a time-consistent reconciliation map, we first construct some valid reconciliation map *μ* from (*T*; *t*, **σ**) to *S*. Using the lca-map ℓ from Algorithm 2, *μ* will be adjusted to become time-consistent, if possible.

By assumption, there is a reconciliation map from (*T*; *t*, **σ**) to *S*. The for-loop (Line (3)-(5)) ensures that each vertex *μ* ∈ *V* obtained a value *μ*(*u*). We continue to show that *μ* is a valid reconciliation map satisfying **(M1)**-**(M3)**.

Assume that *t*(*u*) = ⨀, in this case ℓ(*u*) = **σ**(*u*), and thus **(M1)** is satisfied. If *t*(*u*) = •, it holds that 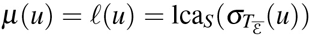, thus satisfying **(M2i)**. Note that ***ρ***_*S*_ ≻_*S*_ ℓ(*u*), and hence, *μ*(*u*) ∈ ε by Line (5), implying that **(M2ii)** is satisfied. Now, assume *t*(*u*) = △ and (*u*, *v*) ∈ ε. By assumption, we know there exists a reconciliation map from *T* to *S*, thus by **(Σ1):**

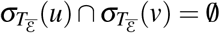

It follows that, ℓ(*u*) is incomparable to ℓ(*v*), satisfying **(M2iii)**.

Now assume that *u*, *v* ∈ *V* and 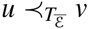. Note that 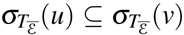. It follows that 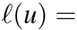 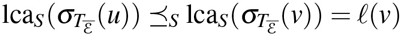. By construction, **(M3)** is satisfied. Thus, *μ* is a valid reconciliation map.

By Theorem 5, two time maps τ_*T*_ and τ_*S*_ satisfying **(D1)**-**(D3)** only exists if the auxiliary graph *A* build on Line (6) is a DAG. Thus if *A*: = *A*_2_ contains a cycle, no such time-maps exists and the statement “No time-consistent reconciliation map exist*S*.” is returned (Line (7)). On the other hand, if *A* is a DAG, the construction in Line (8)-(11) is identical to the construction used in the proof of Theorem 5. Hence correctness of this part of the algorithm follows directly from the proof of Theorem 5.

Finally, we adjust *μ* to become a time-consistent reconciliation map.․ By the latter arguments, τ_*T*_ and τ_*s*_ satisfy **(D1)**-**(D3)** w.r.t. to *μ*. Note, that *μ* is chosen to be the “lowest point” where a vertex *u* ∈ *V* with *t*(*u*) ∈ {◻, △} can be mapped, that is, *μ*(*u*) is set to (*p*(*x*),*x*) where 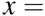 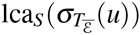. However, by the arguments in the proof of Theorem 2, there is a unique edge (*y*, *z*) ∈ *W* on the path from *x* to ***ρ***_*S*_ such that τ_*s*_(*y*) < τ_*T*_ (*u*) < τ_*s*_(*z*). The latter is ensured by choosing a different value for distinct vertices in *V*(*A*), see comment in Line (9). Hence, Line (14) ensures, that *μ*(*u*) is mapped on the correct edge such that **(C2)** is satisfied. It follows that adjusted *μ* is a valid time-consistent reconciliation map.

We are now concerned with the time-complexity. By Lemma 3, computation of *l* in Line (1) takes *O*(|*V*|log(|*W*|)) time and the for-loop (Line (3)-(5)) takes *O*(<*V*|) time. We continue to show that the auxiliary graph A (Line (6)) can be constructed in *O*(|*V*|log(|*W*|)) time.

Since we know 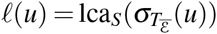 for all *u* ∈ *V* and since *T* and *S* are trees, the subgraph with edges satisfying **(A1)**-**(A3)** can be constructed in *O*(|*V*| + |*W*| + |*E*| + |*F*|) = *O*(|*V*| + |*W*|) time. To ensure **(A4)**, we must compute for a possible transfer edges (*u*, *v*) ∈ ε the vertex 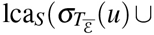 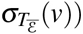. which can be done in *O*(log(|*W*|)) time. Note, the number of transfer edges is bounded by the number of possible transfer event *O*(|*V*|). Hence, generating all edges satisfying **(A4)** takes *O*(|*V*|(log(|*W*|)) time. In summary, computing *A* can done in *O*(|*V*| + |*W*| + |*V*|(log(|*W*|)) = *O*(|*V*|(log(|*W*|)) time.

To detect whether *A* contains cycles one has to determine whether there is a topological order τ on *V*(*A*) which can be done via depth first search in *O*(|*V*(*A*)| + |*E*(*A*)|) time. Since |*V*(*A*)| = |*V*| + |*W*| and *O*(|*E*(*A*)|) = *O*(|*F*| + |*E*| + |*W*| + |*V*|) and *S*, *T* are trees, the latter task can be done in *O*(|*V*| + |*W*|) time. Clearly, Line (10)-(11) can be performed on *O*(|*V*| + |*W*|) time.

Finally, we have to adjust *μ* according to τ_*T*_ and τ_*s*_. Note, that for each *u* ∈ *V* with *t*(*u*) ∈ {◻, △} (Line (12)) we have possibly adjust *μ* to the next edge (*p*(*x*), *x*). However, the possibilities for the choice of (*p*(*x*),*x*) is bounded by by the height of *S*, which is in the worst case log(|*W*|). Hence, the for-loop in Line (12) has total-time complexity *O*(|*V*| log(|*W*|)).

In summary, the overall time complexity of Algorithm 1 is *O*(|*V*| log(|*W*|)).

So far, we have shown how to find a time consistent reconciliation map *μ* given a species tree *S* and a single gene tree *T*. In practical applications, however, one often considers more than one gene family, and thus, a set of gene trees *F* = {(*T*_1_; *t*_1_, **σ**_1_),…, (*T*_*n*_; *t*_*n*_, **σ**_*n*_)} that has to be reconciled with one and the same species tree *S*.

In this case it is possible to aggregate all gene trees (*T*_*i*_; *t*_*i*_ **σ**_*i*_) ∈ *F* to a single gene tree (*T*; *t*, **σ**) that is constructed from *F* by introducing an artificial duplication as the new root of all *T*_*i*_. More precisely, *T* = (*V*, *E*) is constructed from *F* such that 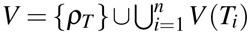 and 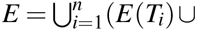 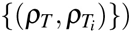. Moreover, the event-labeling map *t* is defined as

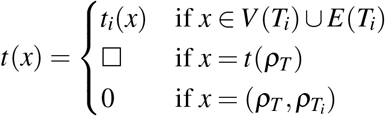

Finally, **σ**(*x*) = *μ*(*x*) for all 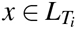.

Finding a time consistent reconciliation for a species tree *S* and a set of gene trees *F* then corresponds to finding a time map τ_*s*_ for *S* and a time map τ_*T*_ for the aggregated gene tree (*T*; *t*, **σ**), such that **(D1)**-**(D3)** are satisfied.

If there exists a time consistent reconciliation map *μ* from (*T*;*t, σ*) to *S* then, by Theorem 2, there exists the two time maps τ_*T*_ and τ_*s*_ that satisfy **(D1)**-**(D3)**. But then τ_*T*_ and τ_*s*_ also satisfy **(D1)**-**(D3)** w.r.t. any (*T*_*i*_; *t*_*i*_, **σ**_*i*_) ∈ *F* and therefore, *μ* immediately gives a time-consistent reconciliation map for each (*T*_*i*_; *t*_*i*_ **σ**_*i*_) ∈ *F*.

## 5 Outlook and Summary

We have characterized here whether a given event-labeled gene tree (*T*; *t*, **σ**) and species tree *S* can be reconciled in a time-consistent manner in terms of two auxiliary graphs *A*_1_ and *A*_2_ that must be DAGs. These are defined in terms of given reconciliation maps. This condition yields an *O*(|*V*| log(|*W*|))-time algorithm to check whether a given reconciliation map *μ* is time-consistent, and an algorithm with the same time complexity for the construction of a time-consistent reconciliation maps, provided one exists.

Our results depend on three conditions on the event-labeled gene trees that are motivated by the fact that event-labels can be assigned to internal vertices of gene trees only if there is observable information on the event. The question which event-labeled gene trees are actually observable given an arbitrary, true evolutionary scenario deserves further investigation in future work. Here we have used conditions that arguable are satisfied when gene trees are inferred using sequence comparison and synteny information. A more formal theory of observability is still missing, however.

Our results point to an efficient way of deciding whether a *given* pair of gene and species tree can be time-consistently reconciled. Such gene and species trees can be obtained from genomic sequence data using the following workflow: (i) Estimate putative orthologs and HGT events using e.g. one of the methods detailed in [3, 1, 4, 8, 31, 32, 34, 39, 41, 43] and [9, 10, 30, 36, 37], respectively. Importantly, this step uses only sequence data as input and does not require the construction of either gene or species trees. (ii) Correct these estimates in order to derive “biologically feasible” homology relations as described in [18, 25, 12, 28, 29, 11, 13, 27, 1]. The result of this step are (not necessarily fully resolved) gene trees together with event-labels. (iii) Extract “informative triples” from the event-labeled gene tree. These imply necessary conditions for gene trees to be biologically feasible [18, 25].

In general, there will be exponentially many putative species trees. This begs the question whether there is *at least one* species tree *S* for a gene tree and if so, how to construct s. In the absence of HGT, the answer is known: time-consistent reconciliation maps are fully characterized in terms of “informative triples” [25]. Hence, the central open problem that needs to be addressed in further research are sufficient conditions for the existence of a time-consistent species tree *given* an event-labeled gene tree with HGT.

## 6 Proof of Theorem 1

We show that Definition 2 is is equivalent to the traditional definition of a DTL-scenario [40, 5] in the special case that both the gene tree and species trees are binary. To this end we establish a series of lemmas detailing some useful properties of reconciliation maps.

### Lemma 4.

*Let μ be a reconciliation map from* (*T*; *t*, **σ**) *to S and assume that T is binary. Then the following conditions are satisfied:*

1. *If v, w* ∈ *V*(*T*) *are in the same connected component of* 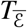, *then* 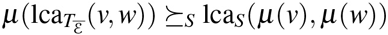.

*Let u be an arbitrary interior vertex of T with children v, w, then:*

1. *μ* (*u*) *and μ* (*v*) *are incomparable in S if and only if* (*u, v*) ∈ ε.
2. *If t* (*u*) = •, *then μ*(*v*) *and μ*(*w*) *are incomparable in S*.
3. *If μ*(*v*),*μ*(*w*) *are comparable or μ*(*u*) ≻_*S*_ lca_*S*_(*μ*(*v*),*μ*(*w*)), *then t*(*u*) = ◻.

*Proof*. We prove the Items 1 - 4 separately. Recall, Lemma 1 implies that 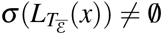 for all *x* ∈ *V*(*T*).

*Proof of Item 1:* Let *v* and *w* be distinct vertices of *T* that are in the same connected component of 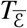. Consider the unique path *P* connecting *w* with *v* in 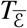. This path *P* is uniquely subdivided into a path *P*′ and a path *P*″ from 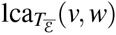 to *v* and *w*, respectively. Condition **(M3)** implies that the images of the vertices of *P*′ and *P*″ under *μ*, resp., are ordered in *S* with regards to ⪯_*S*_ and hence, are contained in the intervals *Q*′ and *Q*″ that connect 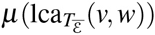 with *μ*(*v*) and *μ*(*w*), respectively. In particular, 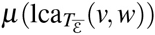 is the largest element (w.r.t. ⪯_*S*_) in the union of *Q*′ ⋃ *Q*″ which contains the unique path from *μ*(*v*) to *μ*(*w*) and hence also lca_*S*_(*μ*(*v*), *μ*(*w*)).

*Proof of Item 2:* If (*u, v*) ∈ ε then, *t*(*u*) = △ and **(M2iii)** implies that *μ*(*u*) and *μ*(*v*) are incomparable.

To see the converse, let *μ*(*u*) and *μ*(*v*) be incomparable in *S*. Item **(M3)** implies that for any edge 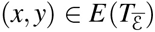 we have *μ*(*y*) ⪯_*S*_ *μ*(*x*). However, since *μ*(*u*) and *μ*(*v*) are incomparable it must hold that 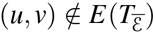. Since (*u, v*) is an edge in the gene tree *T*, (*u, v*) ∈ ε is a transfer edge.

*Proof of Item 3:*

Let *t*(*u*) = •. Since none of (*u, v*) and (*u, w*) are transfer-edges, it follows that both edges are contained in 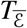.

Then, since *T* is a binary tree, it follows that 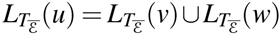 and therefore, 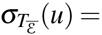 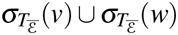.

Therefore and by Item **(M2i)**,

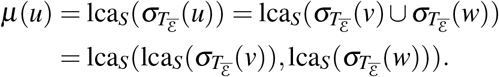

Assume for contradiction that *μ* (*v*) and *μ* (*w*) are comparable, say, *μ*(*w*) ⪰_*S*_ *μ* (*v*). By Lemma 2, 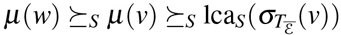 and 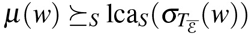. Thus,

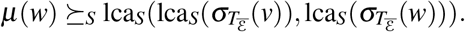

Thus, *μ*(*w*) ⪰_*S*_ *μ*(*u*); a contradiction to **(M3ii)**.

*Proof of Item 4:* Let *μ*(*v*), *μ*(*w*) be comparable in *S*. Item 3 implies that *t*(*u*) ≠ •. Assume for contradiction that *t*(*u*) = △. Since by **(O2)** only one of the edges (*u*, *v*) and (*u, w*) is a transfer edge, we have either (*u*, *v*) ∈ ε or (*u, w*) ∈ ε. W.l.o.g. let (*u,v*) ∈ ε and 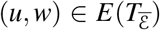. By Condition **(M3)**, *μ*(*u*) ⪰_*S*_ *μ*(*w*). However, since *μ*(*v*) and *μ*(*w*) are comparable in *S*, also *μ*(*u*) and *μ*(*v*) are comparable in *S*; a contradiction to Item 2. Thus, *t*(*u*) ≠ △. Since each interior vertex is labeled with one event, we have *t*(*u*) = ◻.

Assume now that *μ*(*u*) ≻_*S*_ lca_*S*_ (*μ*(*v*), *μ*(*w*)). Hence, *μ*(*u*) is comparable to both *μ*(*v*) and *μ*(*w*) and thus, **(M2iii)** implies that *t*(*u*) ≠ △. Lemma 2 implies 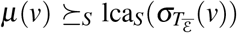 and 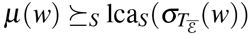. Hence,

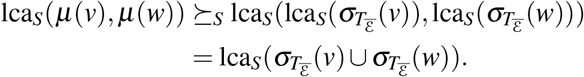

Since *T*(*u*) ≠ △ it follows that neither (*u*, *v*) ∈ ε nor (*u, w*) ∈ ε and hence, both edges are contained in 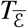. By the same argumentation as in Item 3 it follows that 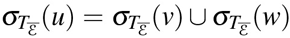 and therefore, 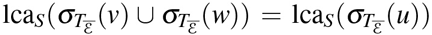. Hence, 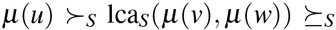 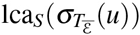. Now, (**M2i**) implies t(*u*) ≠ •. Since each interior vertex is labeled with one event, we have t(*u*) = ◻.

### Lemma 5.

*Let μ be a reconciliation map for the gene tree* (*T*; *t*, **σ**) *and the species tree S as in Definition 2. Moreover assume that T and S are binary. Setfor all u* ∈ *V*(*T*):

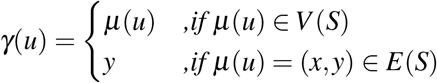

*Then γ*: *V*(*T*) → *V*(*S*) *is a map according to the DTL-scenario.*

*Proof.* We first emphasize that, by construction, *μ*(*u*) ⪰_*S*_ *γ*(*u*) for all u ∈ *V*(*T*). Moreover, *μ*(*u*) = *μ*(*v*) implies that *γ*(*u*) = *γ*(*v*), and *γ*(*u*) = *γ*(*v*) implies that *μ*(*u*) and *μ*(*v*) are comparable. Furthermore, *μ*(*u*) ≺_*S*_ *μ*(*v*) implies *γ*(*u*) ≼_*S*_ *γ*(*v*), while *γ*(*u*) ≺_*S*_ *γ*(*v*) implies that *μ*(*u*) ≺_*S*_ *μ*(*v*).

Thus, *μ*(*u*) and *μ*(*v*) are comparable if and only if *γ*(*u*) and *γ*(*v*) are comparable.

Item (I) and **(M1)** are equivalent.

For Item (II) let *u* ∈ *V*(*T*) \ 𝔾 be an interior vertex with children *v, w*. If (*u, w*) ∉ ε, then 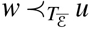. Applying Condition **(M3)** yields *μ*(*w*) ⪯_*S*_ *μ*(*u*) and thus, by construction, *γ*(*w*) ⪯_*S*_ *γ*(*u*). Therefore, *γ*(*u*) is not a proper descendant of *γ*(*w*) and *γ*(*w*) is a descendant of *γ*(*u*). If one of the edges, say (*u*, *v*), is a transfer edge, then *t*(*u*) = △ and by Condition **(M2iii)** *μ*(*u*) and *μ*(*v*) are incomparable. Hence, *γ*(*u*) and *γ*(*v*) are incomparable. Therefore, *γ*(*u*) is no proper descendant of *γ*(*v*). Note that **(O2)** implies that for each vertex *u* ∈ *V*(*T*) \ 𝔾 at least one of its outgoing edges must be a non-transfer edge, which implies that *γ*(*w*) ⪯_*S*_ *γ*(*u*) or *γ*(*v*) ⪯_*S*_ *γ*(*u*) as shown before. Hence, Item (IIa) and (IIb) are satisfied.

For Item (III) assume first that (*u*, *v*) ∈ ε and therefore *t*(*u*) = △. Then, **(M2iii)** implies that *μ*(*u*) and *μ*(*v*) are incomparable and thus, *γ*(*u*) and *γ*(*v*) are incomparable. Now assume that (*u*, *v*) is an edge in the gene tree *T* and *γ*(*u*) and *γ*(*v*) are incomparable. Therefore, *μ*(*u*) and *μ*(*v*) are incomparable. Now, apply Lemma 4(2).

Item (IVa) is clear by the event-labeling *t* of *T* and since **(O2)**. Now assume for (IVb) that *t*(*u*) = •. Lemma 4(3) implies that *μ*(*v*) and *μ*(*w*) are incomparable and thus, *γ*(*v*) and *γ*(*w*) must be incomparable as well. Furthermore, Condition **(M2i)** implies that 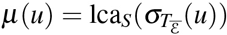). Lemma 2 implies that 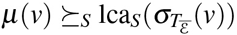 and 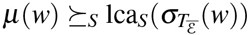. The latter together with the incomparability of *μ*(*v*) and *μ*(*u*) implies that

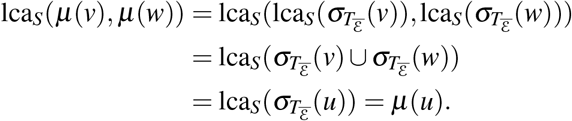

If *μ*(*v*) is mapped on the edge (*x,y*) in *T*, then *γ*(*v*) = *y*. By definition of lca for edges, lca_*S*_(*μ*(*v*), *γ*(*w*)) = lca_*S*_(*y*, *γ*(*w*)) = lca_*S*_(*γ*(*v*), *γ*(*w*)). The same argument applies if *μ*(*w*) is mapped on an edge. Since for all *z* ∈ *V*(*T*) either *μ*(*z*) ≻_*S*_ *γ*(*z*) (if *μ*(*z*) is mapped on an edge) or *μ*(*z*) = *γ*(*z*), we always have

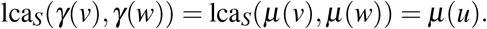

Since *t*(*u*) = •, **(M2i)** implies that *μ*(*u*) ∈ *V*(*S*) and therefore, by construction of *γ* it holds that *μ*(*u*) = *γ*(*u*). Thus, *γ*(*u*) = lca_*S*_(*γ*(*v*), *γ*(*w*)). For (IVc) assume that *t*(*u*) = ◻. Condition **(M3)** implies that *μ*(*u*) ⪰_*S*_ *μ*(*v*), *μ*(*w*) and therefore, *γ*(*u*) ⪰_*S*_ *γ*(*v*), *γ*(*w*). If *γ*(*v*) and *γ*(*w*) are incomparable, then *γ*(*u*) ⪰_*S*_ *γ*(*v*), *γ*(*w*) implies that *γ*(*u*) ⪰_*S*_ lca_*S*_(*γ*(*v*), *γ*(*w*)). If *γ*(*v*) and *γ*(*w*) are comparable, say *γ*(*v*) ⪰_*S*_ *γ*(*w*), then *γ*(*u*) *μ*_*S*_ *γ*(*v*) = lca_*S*_(*γ*(*v*), *γ*(*w*)). Hence, Statement (IVc) is satisfied.

### Lemma 6.

*Let γ*: *V*(*T*) → *V*(*S*) *be a map according to the DTL-scenario for the binary the gene tree* (*T*; *t*, **σ**) *and the binary species tree S. For all u* ∈ *V*(*T*) *define:*

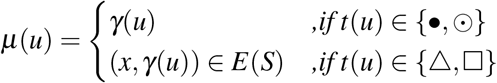

*Then μ*: *V*(*T*) → *V*(*S*) ⋃ *E*(*S*) *is a reconciliation map according to Definition 2*.

*Proof*. Let *γ*: *V*(*T*) → *V*(*S*) be a map a DTL-scenario for the binary the gene tree (*T*; *t*, **σ**) and the species tree *S*.

Condition **(M1)** is equivalent to (I).

For **(M3)** assume that 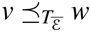. The path *P* from *v* to *w* in 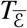 does not contain transfer edges. Thus, by (III) all vertices along *P* are comparable. Moreover, by (IIa) we have that *γ*(*w*) is not a proper descendant of the image of its child in *S*, and therefore, by repeating these arguments along the vertices *x* in *P*_*wv*_, we obtain *γ*(*v*) ⪯_*S*_ *γ*(*x*) ⪯_*S*_ *γ*(*w*).

If *γ*(*v*) ≺_*S*_ *γ*(*w*), then by construction of *μ*, it follows that *μ*(*v*) ≺_*S*_ *μ*(*w*). Thus, **(M3)** is satisfied, whenever *γ*(*v*) ≺_*S*_ *γ*(*w*). Assume now that *γ*(*v*) = *γ*(*w*). If *t*(*v*), *t*(*w*) ∈ {◻, △} then *μ*(*v*) = (*x*, *γ*(*v*)) = (*x*, *γ*(*w*)) = *μ*(*w*) and thus **(M3i)** is satisfied. If *t*(*v*) = • and *t*(*w*) ≠ • then since *μ*(*v*) = *γ*(*v*) and *μ*(*w*) = (*x*, *γ*(*w*)). Thus *μ*(*v*) ≺_*S*_ *μ*(*w*).

Now assume that *γ*(*v*) = *γ*(*w*) and *w* is a speciation vertex. Since *t*(*w*) = •, for its two children w′ and w″ the images *γ*(*w′*) and *γ*(*w″*) must be incomparable due to (IVb). W.l.o.g. assume that w″ is a vertex of *P*_*wv*_. Since *γ*(*V*) ⪯_*S*_ *γ*(*x*) ⪯_*S*_ *γ*(*w*) for any vertex *x* along *P*_*wv*_ and *γ*(*v*) = *γ*(*w*), we obtain *γ*(*w*′) = *γ*(*w*). However, since *γ*(*w*″) ⪯_*S*_ *γ*(*w*), the vertices *γ*(*w′*) and *γ*(*w*″) are comparable in S; contradicting (IVb). Thus, whenever *w* is a speciation vertex, *γ*(*w*′) = *γ*(*w*) is not possible. Therefore, *γ*(*v*) ⪯_*S*_ *γ*(*w*′) ≺_*S*_ *γ*(*w*) and, by construction of *μ*, *μ*(*v*) ≺_*S*_ *μ*(*w*). Thus, **(M3ii)** is satisfied.

Finally, we show that **(M2)** is satisfied. To this end, observe first that **(M2ii)** is fulfilled by construction of *μ* and **(M2iii)** is an immediate consequence of (III). Thus, it remains to show that **(M2i)** is satisfied. Thus, for a given speciation vertex *u* we need to show that 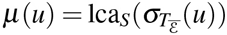.

By construction, *μ*(*u*) = *γ*(*u*). Note, 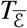 does not contain transfer edges. Applying (III) implies that for all edges (*x*,*γ*) in 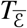 the images *γ*(*x*) and *γ*(*γ*) must be comparable. The latter and (IIa) implies that for all edges (*x*,*γ*) in 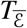 we have *γ*(*y*) ⪯_*S*_ *γ*(*x*). Take the latter together, **σ**(*z*) = *γ*(*z*) ≼_*S*_ *γ*(*u*) for any leaf 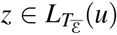. Therefore 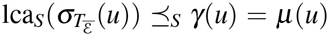. Assume for contradiction that 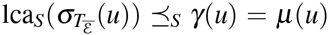. Consider the two children *u*′ and *u*″ of *u* in 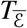. Since neither (*u*, *u*′) ∈ ε nor (*u*, *u*″) ∈ ε and *T* is a binary tree, it follows that 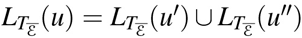 and we obtain that 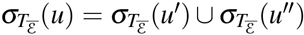. Moreover, re-using the arguments above, 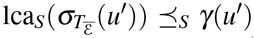 and 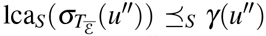. By the arguments we used in the proof for **(M3)**, we have *γ*(*u*′) ≺_*S*_ *γ*(*u*) and *γ*(*u*″) ≺_*S*_ *γ*(*u*). In particular, *γ*(*u*′) and *γ*(*u*″) must be contained in the subtree of *S* that is rooted in the child *a* of *γ*(*u*) in *S* with 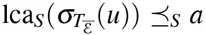, as otherwise, 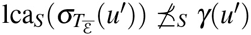 or 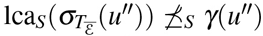. Moreover, neither 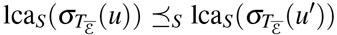 nor 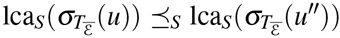 is possible since then 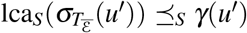 and 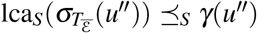 implies that *γ*(*u*′) and *γ*(*u*″) would be comparable; contradicting (IVb). Hence, there remains only one way to locate *γ*(*u*′) and *γ*(*u*″), that is, they must be located in the subtree of *S* that is rooted in 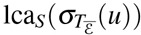. But then we have lca_*S*_(*γ*(*u*′), 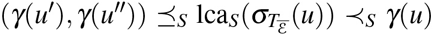; a contradiction to (IVb) *γ*(*u*) = lca_*S*_(*γ*(*u*′), *γ*(*u*″)). Therefore, 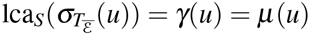 and (**M2i**) is satisfied.

Finally, Lemma 5 and 6 imply Theorem 1.

## Acknowledgment

We thank the organizers of the 32nd TBI Winterseminar 2017 in Bled (Slovenia), where the authors participated, met and basically drafted the main ideas of this paper, while drinking a cold and tasty red Union, or was it a green Laško?

